# Environmental interactions with amoebae as drivers of bacterial-fungal endosymbiosis and pathogenicity

**DOI:** 10.1101/584607

**Authors:** Herbert Itabangi, Poppy C. S. Sephton-Clark, Xin Zhou, Georgina P. Starling, Zamzam Mahamoud, Ignacio Insua, Mark Probert, Joao Correia, Patrick J. Moynihan, Teklegiorgis Gebremariam, Yiyou Gu, Ashraf S. Ibrahim, Gordon D. Brown, Jason S. King, Elizabeth R. Ballou, Kerstin Voelz

## Abstract

Opportunistic infections by environmental fungi are a growing clinical problem, driven by an increasing population of people with immunocompromising conditions. Spores of the Mucorales order are ubiquitious in the environment but can also cause acute invasive infections in humans through germination and evasion of the mammalian host immune system. How they achieve this, and the evolutionary drivers underlying the acquisition of virulence mechanisms, are poorly understood. Here we show that a clinical isolate of *Rhizopus microsporus* contains a *Ralstonia pickettii* bacterial endosymbiont required for virulence in both zebrafish and mice, and that this endosymbiosis enables secretion of factors that potently suppress growth of the soil amoeba *Dictyostelium discoideum*, as well as their ability to engulf and kill other microbes. As amoebae are natural environmental predators of both bacteria and fungi, we propose this tri-kingdom interaction contributes to establishing the endosymbiosis, and acquisition of anti-phagocyte activity. Importantly, we show this activity also protects fungal spores from phagocytosis and clearance by human macrophages, and endosymbiont removal renders the fungal spores avirulent *in vivo*. Together, these findings describe a novel role for a bacterial endosymbiont in *Rhizopus microsporus* pathogenesis in animals, and suggest a mechanism of virulence acquisition through environmental interactions with amoebae.

**In brief:** How environmental fungi evolved the mechanisms that enable them to cause opportunistic infections in humans is unclear. Here, we identify a novel tri-kingdom interaction, whereby a bacterial endosymbiont, living within a clinical isolate of the ubiquitous environmental fungus *Rhizopus microsporus*, causes the generation of a secreted activity that blocks the growth and predatory activity of amoebae. We suggest this provides a new evolutionary driver for the establishment of bacterial/fungal endosymbiosis and demonstrate this is critical for fungal pathogenicity *in vivo*.

## Introduction

Soil-dwelling fungi must evade predation by phagocytic amoebae and, similarly, pathogenic fungi evade host phagocytic cells that defend against infection (1, 2). Interaction with amoebae has been proposed as an evolutionary training ground for pathogenic fungi, enabling them to resist the multifactorial stresses of the mammalian host. Mucoralean fungi such as soil-associated *Rhizopus microsporus*, which causes invasive mucormycosis, successfully establish in complex polymicrobial environments that include predatory amoeba. Yet, they lack common strategies fungi employ to evade predation and phagocytosis: *R. microsporus* fails to mask cell wall ligands in swollen spores, has a slow germination rate, and produces relatively low biomass (3, 4). Mucorales resting spores also fail to elicit pro-inflammatory cytokine responses and do not induce strong phagocyte chemotaxis. Here, we investigate an as yet unexplored aspect of *R. microsporus* pathogen biology: the impact of a bacterial endosymbiont on interaction with amoebae and host phagocytes.

Pathogenic Mucorales span multiple genera, with the *Rhizopus* genus causing almost half of all documented cases and high mortality in susceptible patient populations (5–12). Infection occurs through inoculation with dormant, immunologically inert spores. Upon germination, these spores become metabolically active and begin to swell (13, 14). Based on analogy to *Aspergillus* spores, frequently used as a model for Mucorales, swelling is expected to reveal Microbe Associated Molecular Patterns (MAMPs) and induce increasing rates of phagocytosis which must be overcome by rapid hyphal extension (15, 16). However, swollen spores in the *Rhizopus* genus are often no more readily phagocytosed than resting spores and *R. microsporus* spores are phagocytosed at lower rates than other well studied fungal spores (17–20). While in some species, larger spore size (12.3 μm) reduces phagocytosis, the small spore size of *R. microscoprus* (5 μm, comparable to other well-phagocytosed particles) makes this unlikely to explain the observed reduced uptake (21, 22).

A common strategy fungi employ to evade phagocytosis is through masking cell wall ligands (16, 23–28). Two classic examples of this are the *Cryptococcus neoformans* capsule and the *Aspergillus* spore hydrophobin layer, which effectively block phagocytosis by preventing detection of MAMPs (15, 16, 27–30). However, there is no evidence of hydrophobins in the *R. microsporus* genome, and evidence of cell wall remodelling that masks MAMPs is limited to the hyphal phase (31). Moreover, *R. microsporus* has a slow germination rate, and produces relatively low biomass, which are predicted to reduce fungal virulence (3, 4, 21, 32–34). Together, these differences suggest an alternate strategy available to *Rhizopus* for the evasion of phagocytosis during the early stages of germination.

Mucormycete plant pathogens, including the Mucorales, are frequently colonized or influenced by bacteria of the genera *Burkholderia, Paraburholderia*, and *Ralstonia*, all closely related to the *Pseudomonas* genus (35–37). During endosymbiosis, *Burkholderia* spp. influence fungal behaviour such as growth and sporulation, and also mediate plant pathogenesis (37, 38). For example, the endotoxin Rhizoxin is secreted by *Burkholderia rhizoxina* during mutualism with *R. microsporus* and inhibits plant defences (39, 40). The *Ralstonia solanacearum* lipopeptide ralsolamycin can influence chlamydospore development and hyphal invasion (37). While endosymbionts are widespread in patient fungal samples, a role for endosymbionts in preventing fungal phagocytosis has not been established (35). Rather, work examining this interaction in the context of phagocyte-deficient mouse models found no correlation between endosymbiont presence and fungal virulence (35). We recently showed that phagocyte activity in the early stages of infection control are critical for predicting Mucorales disease outcome, suggesting that phagocyte-deficiency primes the host for infection (41). Together, this raises the hypothesis: do endosymbionts specifically impact fungal interaction with phagocytic cells, including both environmental amoebae and mammalian macrophages and neutrophils, leading to enhanced fungal virulence?

Here, we investigate the interaction of environmental and early host phagocytic cells with *R. microsporus*. We report, for the first time, a role for a bacterial endosymbiont of *R. microsporus* in modulating the interaction with these phagocytic cells and further expand this to demonstrate a role in the early stages of disease in both zebrafish and murine models of infection. Specifically, we observed a significant reduction in phagocytosis of metabolically activated spores compared to resting spores. We investigate the consequences of endosymbiont status on infection outcome and demonstrate that this bacterial endosymbiont contributes to both fungal stress resistance and immune evasion during the earliest stages of infection, enabling fungal pathogenesis.

## Results

### A clinical isolate of *R. microsporus* suppresses phagocytosis by macrophages

We previously showed that the early stages of host-fungus interaction determine disease outcome in the zebrafish model of mucormycosis (41). Successful control of infection therefore requires both the presence of phagocytes at the site of infection within the first 24 hours and their subsequent ability to kill spores. We hypothesized that, in instances where infection control fails, spores might evade phagocytosis. We therefore examined in detail the interactions between spores from *R. microsporus* strain FP469-12, a clinical isolate from a patient at the Queen Elizabeth Hospital, Birmingham, and J774A.1 macrophage-like cells.

During these critical early stages of infection, *R. microsporus* spores become metabolically active and start to swell, prior to germination (42, 43). This swelling is normally associated with exposure of surface MAMPs which should facilitate phagocytosis (1). Contrary to this expectation however, whilst dormant *Rhizopus* spores were readily engulfed by J774A.1 cells, swollen spores were taken up significantly less (**Figure 1A**, p>0.0001). This was dependent on fungal viability as UV-killing of swollen spores completely restored their uptake (**Supp Figure 1A**). Larger objects are harder for phagocytes to engulf, and under these conditions swollen spores reached a mean diameter of 7.3 μm after 6 hours, compared to 4.6 μm during dormancy. However, as J774A.1 cells engulfed latex beads up to 11.9 μm in diameter at least as well as dormant spores (**Supp Figure 1B**), size was not a limiting factor.

**Figure 1:**
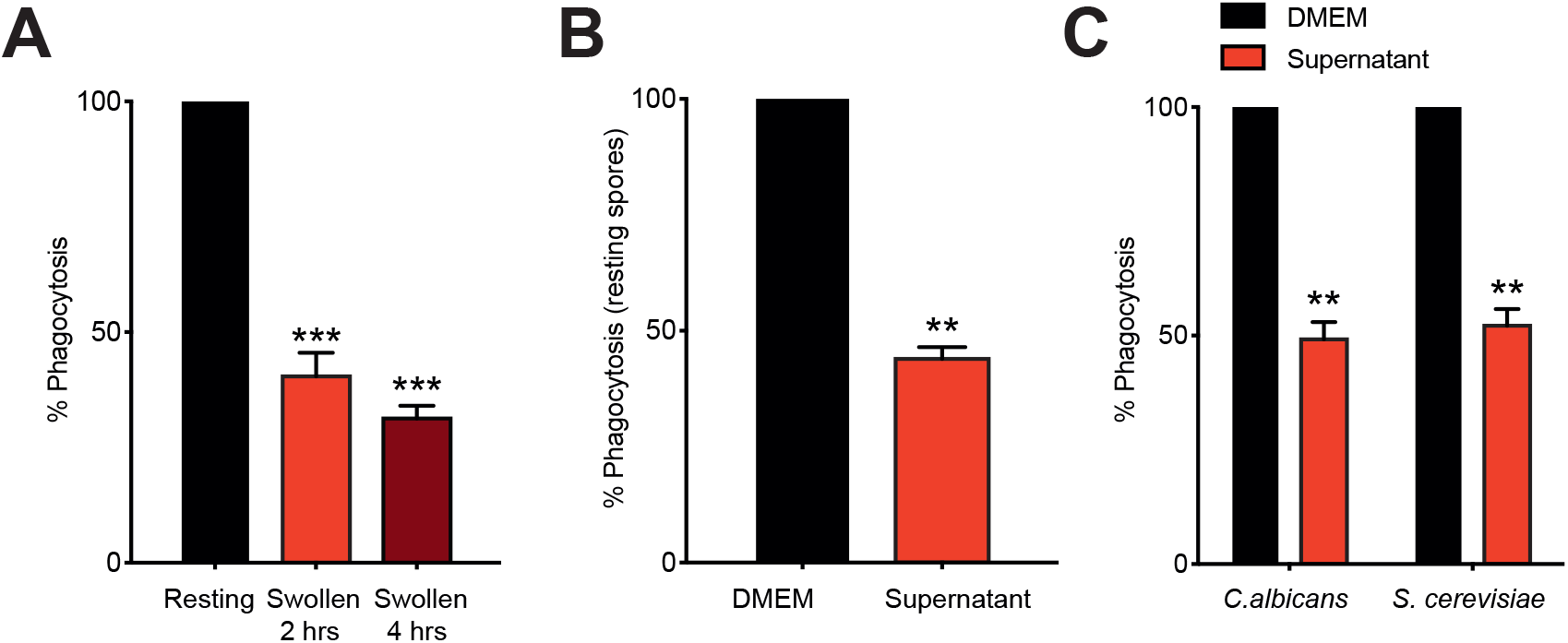
Swelling of *R. microsporus* FP469-12 spores inhibits phagocytosis. (A) Phagocytosis of resting spores, or those allowed to swell for 2 or 4h by J447.2 macrophages. (B) Effect of swollen spore supernatant on phagocytic uptake of naïve, resting *R. microsporus* spores by macrophages. (C) Effect of *R. microsporus* FP469-12 conditioned medium on phagocytosis of *C. albicans* and *S. cerevisiae*. For all assays, the number of macrophages containing at least one spore were counted after 1 hour. n=3 biological replicates of >1000 macrophages each, error bars represent SEM. * = p<0.05, ** p<0.001,*** = p<0.0001, One way ANOVA with Tukey’s correction for multiple comparisons.

As activation of phagocytic receptors by fungal surface ligands is critical for engulfment, we measured how the fungal cell wall changed upon swelling. Whilst total chitin (stained with calcofluor white (CFW), Median Fluorescence Intensity) increased more than 2-fold (**Supp Figure 1C and D**), surface exposure of chitin (WGA) and total protein (FITC) was unaltered between resting and swollen spores (**Supp Fig 1E and F**). We were unable to detect any β-glucan exposure, consistent with previous reports showing a lack of glucan in *Rhizopus* species spore cell walls (3, 44). Instead, mannan (stained with the lectin Concavalin A) was detected as a major component of the cell wall at all stages of growth: exposed mannan was detected on the surface of all cells and was particularly high in approximate half of resting spores and swollen (**Supp Figure 1C and G**). Whilst we cannot exclude differential exposure of additional unknown surface components, in general, cell wall remodelling does not correlate with the observed protection from phagocytosis.

These findings indicate a novel active mechanism for evading phagocytosis upon spore germination. As this effect does not appear to correlate with large-scale changes in fungal cell wall, we hypothesised a secreted factor might be responsible. We therefore allowed *Rhizopus* spores to swell in macrophage medium (DMEM) for 1 hr before removing the spores and testing the capacity of the conditioned medium supernatant to inhibit phagocytosis of other particles. This conditioned medium was sufficient to inhibit phagocytosis of dormant spores to a similar extent as swollen spores themselves **(Figure 1B)**. Conditioned media also had a cross-protective effect on non-mucormycetes, inhibiting phagocytosis of the ascomycete yeasts *Candida albicans* and *Saccharomyces cerevisiae* (**Figure 1C**). *R. microsporus* FP469-12 therefore produces a secreted factor (or factors), induced upon spore swelling and metabolic activation, with broad anti-phagocytic activity.

### *R. microsporus* FP469 harbours a bacterial endosymbiont

Mucorales species are widely associated with bacterial endosymbionts, which can contribute to the synthesis of secreted metabolites (35, 45). The endosymbiont *Burkholderia rhizoxinia* augments fungal pathogenesis of *R. microsporus* in rice seedlings by producing the secondary metabolite Rhizoxin, a toxin that targets a conserved residue in β-tubulin, potently depolymerising microtubules (39, 46–48). However, no involvement for endosymbionts in mammalian disease has been proven, either as a requirement for pathogenesis in humans or associated with disease severity in diabetic mice (35, 45).

To test whether an endosymbiotic bacterial product might influence phagocytosis, we tested for the presence of bacterial endosymbiont 16S rRNA in our *Rhizopus* strain, *R. microsporus* FP469-12, by PCR (**Figure 2A**). *R. microsporus* FP469-12 was positive for 16S rRNA (lane 1), which was lost upon treatment of the fungus with the antibiotic ciprofloxacin (lane 2). Through enzymatic and physical disruption of the fungal cell wall, we were able to isolate endosymbionts from FP469-12, and 16S rRNA could be amplified from the isolated bacteria (lane 3). Moreover, the endosymbiont could be visualized inside *R. microsporus* FP469-12 hyphae using the nucleic acid-binding dye SYTO9, and could be cleared by treatment with ciprofloxacin (**Figure 2B**). For the remainder of this work, unless otherwise stated, spores treated with ciprofloxacin were germinated, passaged through sporulation twice, and frozen down for stocks before use as endosymbiont-free spores to limit the possibility that ciprofloxacin was effecting outcomes.

**Figure 2:**
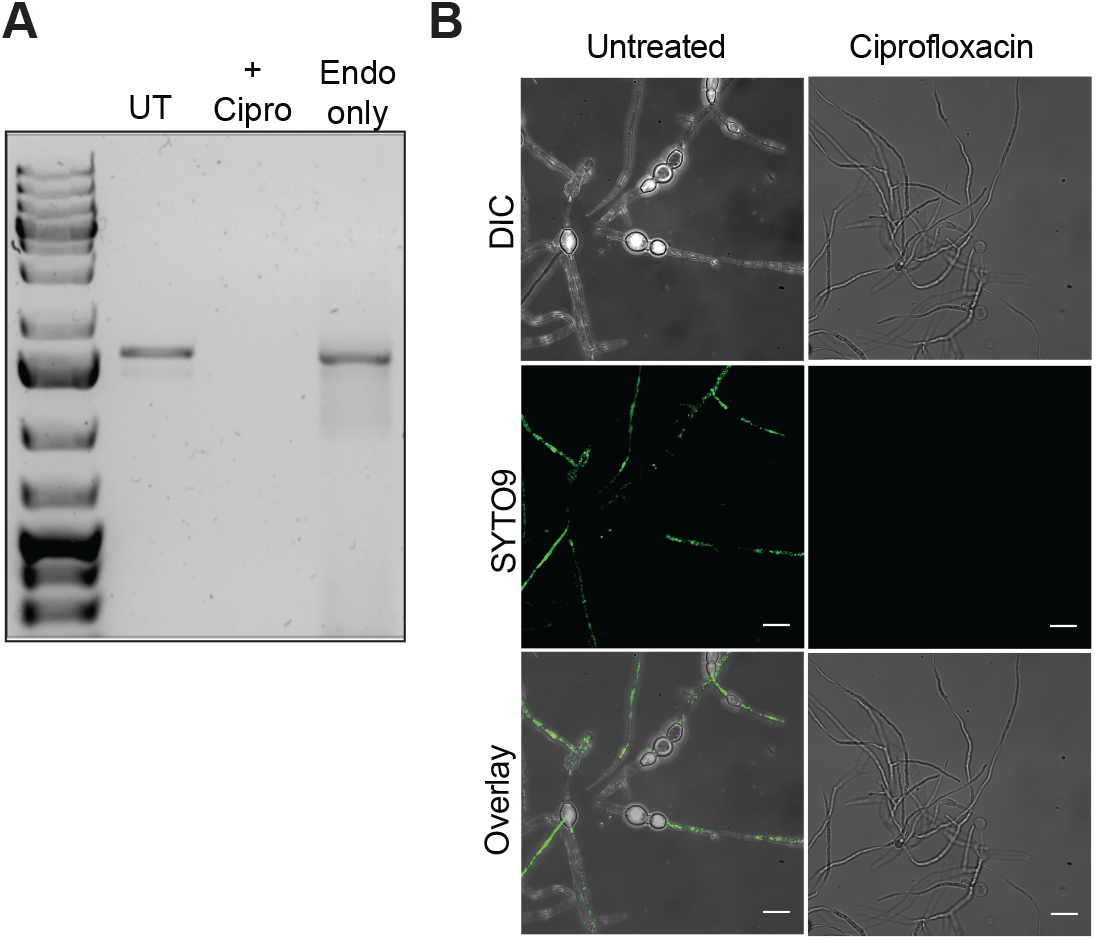
*R. microsporus* FP469-12 contains a bacterial endosymbiont. (A) PCR screen for the presence of bacterial 16S rDNA. Genomic DNA was isolated from wild-type and ciprofloxacin-treated *R. microsporus* FP469-12 cells, as well as the isolated endosymbiont alone. Presence of 16S rDNA is indicated by the presence of a 1.5 kb PCR product. (B) SYTO9 staining for bacterial endosymbionts. Spores of parent and ciprofloxacin-treated R. microsporus FP469-12 were fermented in VK medium, the mycelial pellet submerged in NaCl and then stained with SYTO9 prior to brightfield and fluorescence imaging.

To further characterise the endosymbiont isolated from *R. microsporus* FP469-12, it was subjected to whole genome sequencing. This identified *Ralstonia pickettii*, a relative of *Burkholderia* commonly found in soil and water and occasionally associated with contaminated medical equipment as an opportunistic pathogen (49–52). This was confirmed by two independent replicates, identifying 16S sequences consistent with *R. pickettii* in FP469-12 but not FP469-12 cured genomic extracts.

### Endosymbiotic *R. pickettii* are required for anti-phagocytic activity

We examined whether the presence of the bacterial endosymbiont was necessary for inhibition of fungal spore uptake by macrophages. Uptake of cured swollen spores of *R. microsporus* FP469-12 was significantly higher than that of the uncured parent (p<0.0001) (**Figure 3A**). Media conditioned by endosymbiont-free spores also lost the ability to inhibit phagocytosis (**Figure 3B**). In addition, whilst the *R. pickettii* endosymbiont grew well in the absence of the fungi, it only caused a small decrease in phagocytosis when used to condition macrophage medium, although still significantly more than medium conditioned by cured spores (p<0.001, **Figure 3B**). Endosymbiotic bacteria are therefore required for the secreted anti-phagocytic activity.

**Figure 3:**
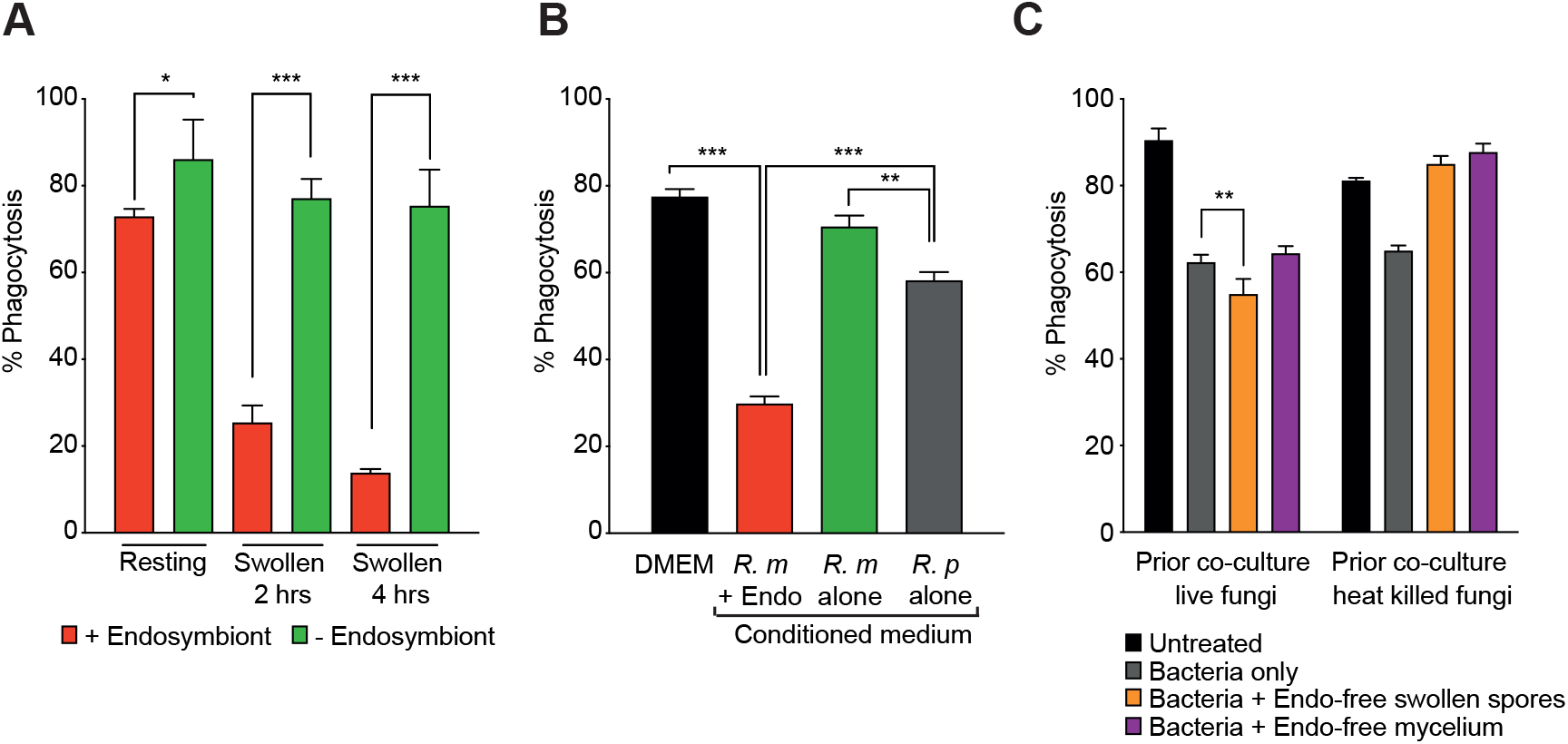
The presence of the endosymbiont is required for anti-phagocytic activity. (A) Phagocytosis of parental, and cirprofloxacin-teated (endosymbiont-free) *R. microsporus* FP469-12 spores by J774.2 macrophages, upon swelling. (B) Contributions of the fungi and bacteria to the secreted anti-phagocytic activity. J774.2 cells were incubated for 1 hr with resting spores in medium conditioned by either parental spores (with endosymbionts), endosymbiont-free spores, or the isolated *R. pickettii* endosymbiont alone. (C) Effect of media conditioned by co-cultures of bacterial symbionts and endosymbiont-free fungal spores grown on phagocytosis of *R. microsporus* resting spores. Each graph shows the mean and SEM of 3 independent experiments. * = p<0.05, **=p<0.001, *** = p<0.000, One-way ANOVA with Tukey’s correction for multiple comparisons.

We next assessed the individual contributions of bacteria and fungus to inhibiting phagocytosis of resting cured fungal spores (**Figure 3B**). There was a small but not significant decrease in phagocytosis when the conditioned media from the fungus alone was used. Conditioned media from bacteria alone caused a decrease in phagocytosis relative to conditioned media from the fungus alone although not as much as when endosymbiotic (**Figure 3B**). We therefore investigated whether the activities were synergistic (**Figure 3C**). Conditioned media generated by co-culturing bacteria with endosymbiont-free swollen spores caused a minor but significant (p=0.019) augmentation of the inhibitory effect of *R. pickettii-*conditioned medium (**Figure 3C**). This effect was lost when either fungal mycelium, or heat-killed spores were used for conditioned medium (**Figure 3C**). Whilst this indicates that both organisms cause a minor additive affect when cultured independently, it never reached the extent seen when growing as a true endosymbiosis. We therefore conclude that fungal/bacterial endosymbiosis is required for effective anti-phagocyte activity.

### Endosymbiosis blocks growth of predatory amoebae

Whilst murcomyetes can cause opportunistic infections in susceptible humans, they normally live in environments such as soil. The emergence of virulence is consequently driven by environmental interactions, rather than those between fungi and the human immune system. It has thus been proposed that mechanisms allowing evasion of phagocytic immune cells originally evolved to help fungi escape professional phagocytes in the environment, such as predatory amoebae (53). To test whether *R. microsporus* FP469-12 can also inhibit capture by environmental phagocytes, we examined its interactions with the soil amoeba *Dictyostelium discoideum*, a well-established model host for both pathogenic bacteria and fungi (54–58).

Consistent with our macrophage data, conditioning *D. discoideum* growth medium (HL5) with swollen *R. microsporus* FP469-12 spores for 4 hours reduced the phagocytosis of inert, heat-killed *S. cerevisiae* by 41%, compared to untreated medium (**Figure 4A**). Timelapse microscopy after addition of conditioned medium revealed *D. discoideum* cells were still able to form protrusions and actively migrate over this period, indicating they remain active and viable. However, the amoebae immediately started accumulating prominent swollen vacuoles (**Figure 4B and Movie 1**). Factors secreted by *R. microsporus* FP469-12 therefore also affect amoebae and inhibit phagocytosis across evolution.

**Figure 4:**
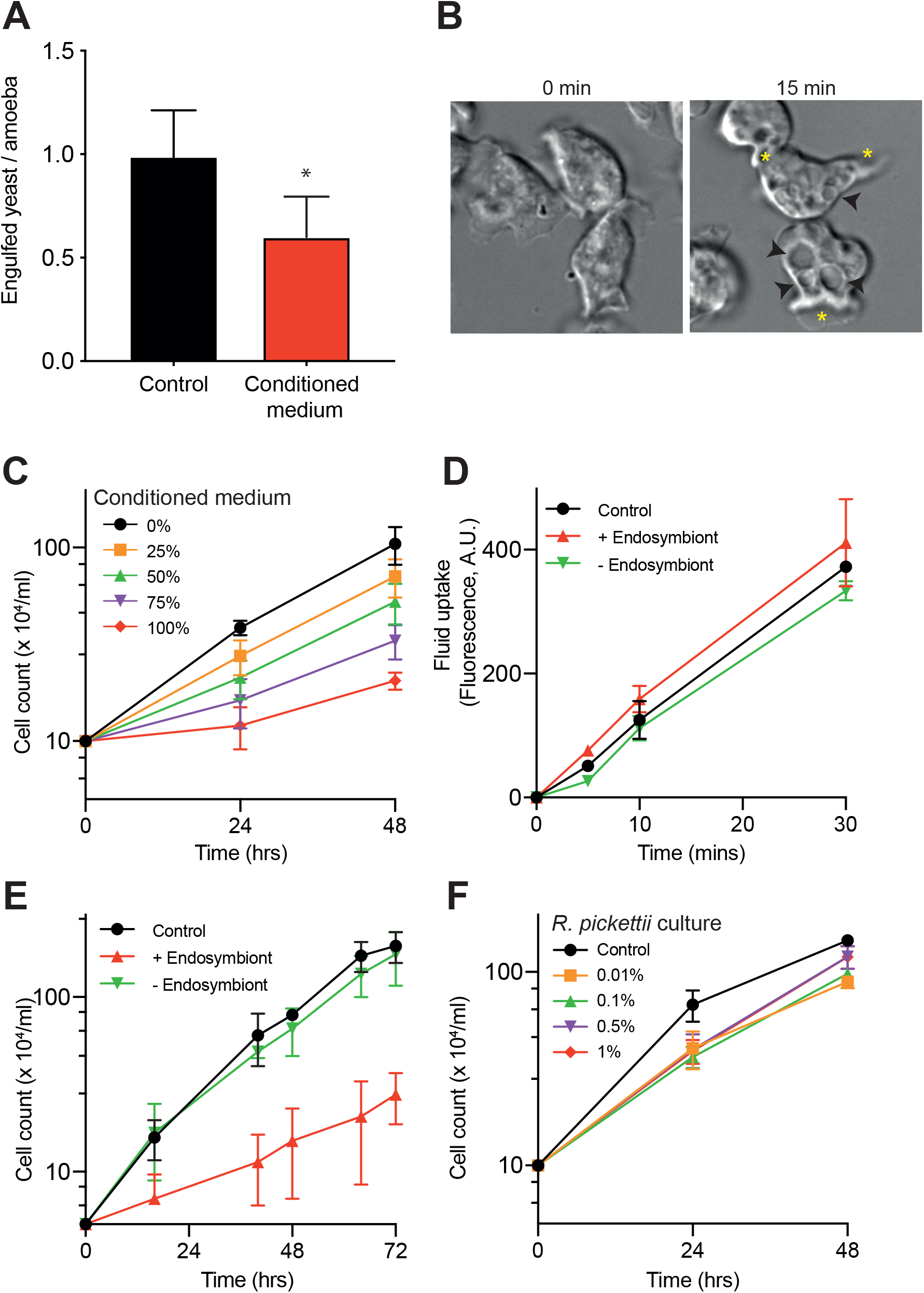
Fungal-bacterial endosymbiosis inhibits growth of amoebae. (A) Phagocytosis of heat-killed *S. cerevisiae* by *D. discoideum* in either normal medium or medium pre-conditioned by *R. microsporus* FP469-12. (B) DIC images of *D. discoideum* cells 0 and 15 minutes after addition of conditioned medium. Black arrowheads indicate the large swollen vacuoles induced, yellow asterisks mark forming protrusions. (C) Dose-dependent inhibition of *D. discoideum* growth by *R. microsporus* FP469-12 conditioned medium. Amoebae were incubated in different concentrations of conditioned medium diluted in fresh medium as indicated, and cells counted at each time point. (D) Fluid uptake (macropinocytosis) by *D. discoideum* cells in medium conditioned by either parental, or endosymbiont-free *R. microsporus* FP469-12 spores. Cells were incubated in TRITC-dextran containing medium, and fluorescent dye uptake measured by flow cytometry. (E) Effect of endosymbiont removal on the ability of *R. microsporus* FP469-12 conditioned medium to inhibit *D. discoideum* growth. (F) Effect of *R. pickettii*-conditioned medium on *D. discoideum* growth. HL5 medium was conditioned for 4 hours by addition of the indicated dilutions of an overnight *R. pickettii* culture, before bacteria were removed. Each graph shows the mean and SEM of 3 independent experiments. * = p<0.05, paired T-test.

The accumulation of swollen vacuoles indicates that *R. microsporus* FP469-12 may cause additional disruption of intracellular vesicle trafficking. Consistent with this, *R. microsporus* FP469-12 conditioned medium caused a strong, dose-dependent inhibition of *D. discoideum* growth (**Figure 4C**). In liquid culture, *D. discoideum* grow using macropinocytosis to take up nutrients. We therefore measured macropinocytosis by following uptake of TRITC-dextran by flow cytometry (**Figure 4D**). This was unaffected by *R. microsporus* FP469-12 conditioned media, indicating defective nutrient uptake was not the cause of inhibited growth. However, growth inhibition was completely dependent on the presence of the endosymbiont, as neither ciprofloxacin treated spores nor ciprofloxacin alone affected *D. discoideum* growth (**Figure 4E**). HL5 medium conditioned with concentrations of the *R. pickettii* endosymbiont well exceeding that present in the endosymbiotic cultures also had little effect on *D. discoideum* growth (**Figure 4F**). Both bacteria and fungi are therefore essential for inhibition of amoeba growth.

### *R. microsporus/ R. pickettii* endosymbiosis inhibits phagosomal maturation and killing

Amoebae obtain nutrients for growth through capture and degradation of microbes or extracellular fluid in phagosomes or macropinosomes respectively (59). As this represents a major evolutionary interface in the competition between microbial predators and prey, we tested whether *R. microsporus* FP469-12 affected *D. discoideum* phagosomal maturation. Using DQ-BSA coated beads, which increase fluorescence upon proteolysis, we found addition of spore-conditioned medium after phagocytosis severely inhibited degradation (**Figure 5A**); this was again dependent on the presence of the endosymbiont. Pre-treating cells for 30 minutes prior to phagocytosis had no additional effect, indicating that inhibition of proteolysis was due to direct interference with the proteolytic machinery, rather than initial lysosomal delivery to the phagosome (**Supp Figure 2A**).

**Figure 5:**
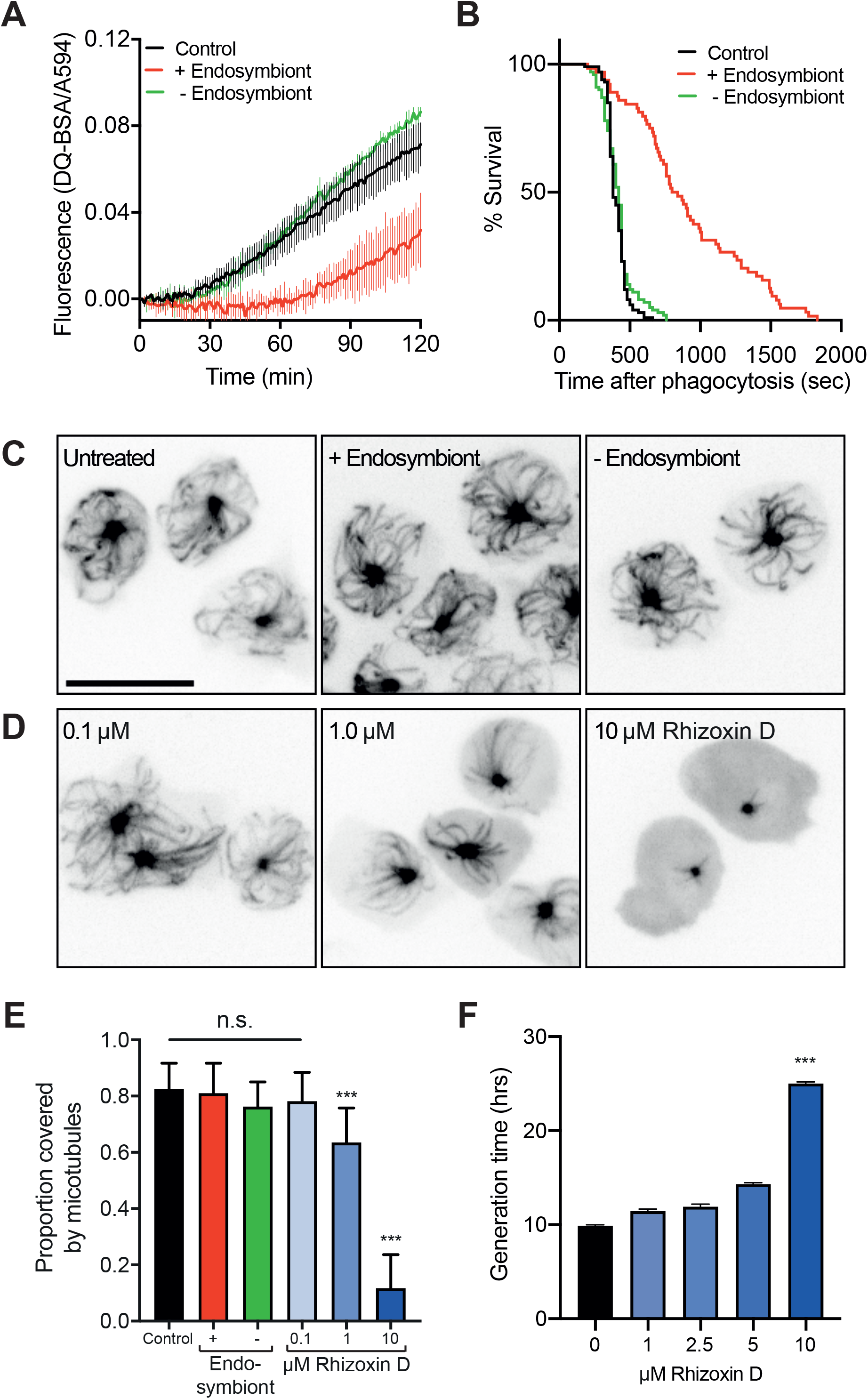
The secreted activity inhibits phagosome maturation by a novel mechanism. (A) Phagosomal proteolysis of *D. discoideum* cells incubated in fungal-conditioned medium, in the presence, or absence of the endosymbiont. Measured by increasing fluorescence of DQ-BSA-conjugated beads after engulfment. (B) Kaplan-Meijer survival curve of GFP-expressing *K. pneumonia* after engulfment by *D. discoideum* in *R. microsporus* FP469-12-conditioned medium. Phagocytosis was observed by timelapse fluorescence microscopy, and the point of bacterial death inferred from the quenching of GFP fluorescence (n>100 for each condition, *** P<0.001, log-rank Mantel-Cox test). (C) Maximum intensity projections of *D. discoideum* cells expressing GFP-α tubulin after 20 minutes treatment with either conditioned medium or (D) the indicated concentrations of Rhizoxin D. (E) Quantification of the proportion of cytoplasm covered by the microtubule array in cells treated as in (C) and (D)(***P<0.001, T-test). (F) Effect of rhizoxin D treatment on *D. discoideum* growth. Generation times calculated from growth curves obtained over 72 hours (***P<0.001, paired T-test). Unless otherwise indicated all graphs show the mean and standard deviations of 3 independent experiments.

We also measured whether *R. microsporus* FP469-12-conditioned medium inhibited the ability of amoebae to kill phagocytosed bacteria. Following the phagocytosis of non-pathogenic *Klebsiella pneumoniae* expressing GFP by timelapse microscopy allows their intracellular survival to be measured, as the GFP-fluorescence becomes quenched upon death and permeabilization in acidic phagosomes (60). In untreated medium, *D. discoideum* killed 50% of engulfed bacteria within 400 seconds. Bacterial survival was significantly promoted by conditioned medium, but only in the presence of the endosymbiont (**Figure 5B**). As amoebae normally feed on bacteria such as *R. pickettii* in the soil, we propose this constitutes an important evolutionary driver of endosymbiosis, whereby collaboration between the bacteria and fungi can inhibit both uptake and killing by amoeboid predators.

### Amoebae are inhibited by a novel, microtubule-independent activity

Both endosymbiosis and the secretion of bioactive metabolites are relatively common in fungi. The previously described *B. rhizoxina* endosymbiont synthesises the rhizoxin family of secondary metabolites via a *trans-*Acyl Transferase Non-Ribosomal Peptide Synthase Polyketide Synthase (*trans-*AT NRPS PKS) gene cluster (39, 61). Rhizoxin directly binds the β-tubulin subunit of microtubules, potently causing their depolymerisation and mitotic arrest in both plants and humans (46). It is thought that this endosymbiosis emerged to help *R. microsporus* damage and saprophytically feed on plant tissue (62). Genomic analysis of the *R. pickettii* isolated from *R. microsporus* FP469-12 using antiSMASH (5 beta, (63)) software for secondary metabolite clusters did not identify a *trans-*AT PKS, although it did reveal a Type I Polyketide Synthase (Type I PKS), as well as biosynthetic pathways for siderophores, bacteriocins, terpenes, and arylpolyenes, among others. The microtubule cytoskeleton is important for many cellular functions, including vesicle trafficking, which is important for phagosome maturation. Therefore, whilst the *R. pickettii* endosymbiont is unlikely to synthesise rhizoxin, we investigated whether microtubules were a general target for *Rhizopus* endosymbionts.

Contrary to this hypothesis, treatment of *D. discoideum* cells expressing GFP-α-tubulin with *R. microsporus* FP469-12 conditioned medium caused no measurable depolymerisation of the microtubule cytoskeleton either with, or without, the endosymbiont (**Figure 5C**). This was quantified by the extent of the microtubule array as a proportion of the cell area (**Figure 5E**). As a positive control, we also treated the cells with rhizoxin D. This was highly effective at depolymerising microtubules, but only at concentrations above 1μM (**Figure 5D and E**). This is 5 orders of magnitude higher than that required for a similar effect in mammalian cells (64), even though all residues at the β-tubulin rhizoxin binding site are conserved (46). Furthermore, whilst conditioned medium blocks *D. discoideum* growth without obvious microtubule disruption, rhizoxin D only inhibited growth at concentrations where microtubules were almost completely depolymerised (**Figure 5F**). We therefore conclude that the *R. microsporus/ R. pickettii* endosymbiosis inhibits amoeba function by a novel, most likely microtubule-independent, mechanism.

### Endosymbiosis facilitates fungal evasion of macrophages

The data above show that endosymbiosis allows *R. pickettii* and *R. microsporus* FP469-12 to inhibit both phagocytosis and phagosomal killing by amoebae. To test whether this promotes virulence in animals, we first tested how the presence of the endosymbiont effected evasion from J774A.1 macrophages. Co-incubation showed that spores containing endosymbiont were highly resistant to clearance by macrophages, with no significant decrease in CFU over 24 hours. In contrast, spores lacking endosymbionts were cleared much more effectively, with 60% removed within 24 hours (**Figure 6A**, p<0.0001).

**Figure 6:**
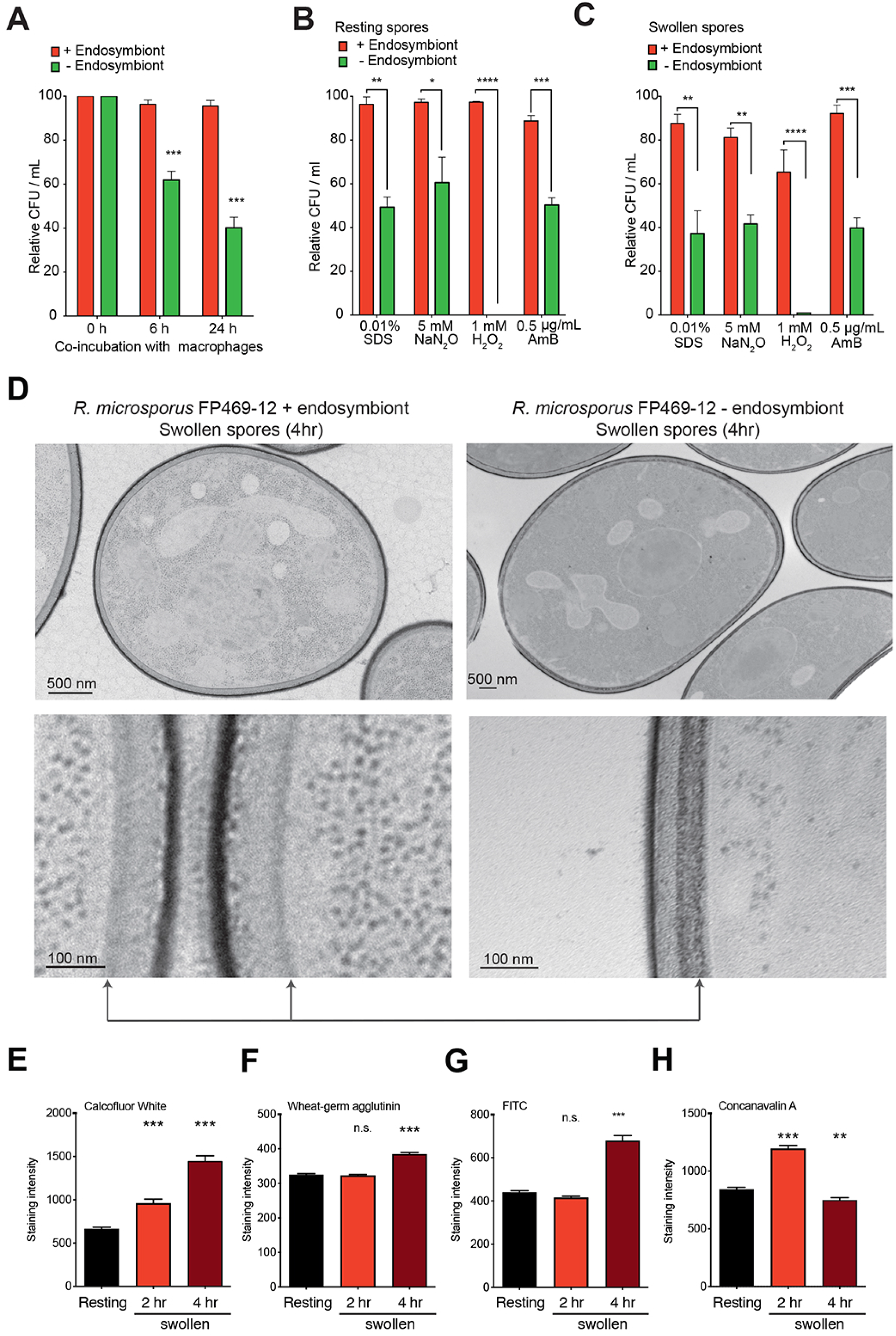
Endosymbiosis with *R. pickettii* influences the fungal cell wall and stress tolerance. (A) Survival (CFUs) of *R. microsporus* FP469-12 resting spores after incubation with J774.1 macrophages in the presence and absence of endosymbiont. (B) and (C) Impact of the endosymbiont on *R. microsporus* FP469-12 spore survival under cell wall, nitrosative, oxidative and antifungal stresses. Either resting (B) or swollen (C) spores were incubated at the indicated concentrations for 24 hours, prior to CFU determination. Graphs (A-C) show mean ± SEM of 3 repeats (**=p<0.001, ***=p=0.0001, ****=p<0.00001, One way ANOVA with Tukey’s correction for multiple comparisons). (D) Shows TEM images of spores swollen for 4 hours in the presence or absence of the endosymbiont. Lower panels show enlargements of representative cell wall regions. (E-H) Show changes in cell wall composition upon swelling of endosymbiont-free fungal spores. Staining intensities were quantified by fluorescence microscopy. Comparable measurements of the parental strain are shown in Supplemental Figure 1D-G. (E) Shows total chitin (Calcofluor White), (F) exposed chitin (Wheat-Germ Agglutinin), (G) total protein (FITC) and (H) mannan (Concanavalin A) (n=300 for each).

The secreted activity identified above may not be the sole protective mechanism conferred by the endosymbiont. Previous work has shown that endosymbiosis with *B. rhizoxina* is also beneficial for fungal development, cell wall synthesis and stress resistance (65). As the fungal cell wall is the main interface between the engulfed fungi and the antimicrobial activities of the phagosome, we investigated whether *R. pickettii* endosymbiosis also affects *R. microsporus* resistance to phagosome-relevant stresses. In the absence of the endosymbiont, both resting and swollen spores were significantly more sensitive to treatment with 0.01% SDS, 5 mM NaNO_3_, or 1 mM H_2_O_2_, as well the front-line antifungal Amphotericin B (0.5 μg/ml) (**Figure 6B and C**). Endosymbiosis with *R. pickettii* therefore increases the resistance of *R. microsporus* to several physiologically important antimicrobial stresses.

We also examined the effect of endosymbiosis on the fungal cell wall structure. TEM of swollen spores with and without the endosymbiont revealed differences in electron density and width of the cell wall interior, suggesting the endosymbiont influences spore cell wall organization (**Figure 6D**). In the presence of the endosymbiont, the only changes we detected upon spore swelling were an increase in polysaccharide levels over time whilst total protein, exposed chitin and mannan levels remained static (**Supp. Figure 1D-G**). Polysaccharide levels still increased when the endosymbiont was removed, although there was less calcofluor white staining at each timepoint compared to the parental strain (**Figure 6E and Supp Figure 1D**). In contrast to the parental strain, both exposed chitin (WGA) and total protein (FITC) levels increased after 4 hours swelling in the absence of the endosymbiont (**Figure 6F and G**) as well as a transient increase in mannan (ConA) at 2 hours swelling (**Figure 6H**, p<0.0001). The presence of the endosymbiont therefore has broad influence on the structure of the cell wall, during swelling and protects both the fungus and bacteria from environmental phagocytes and immune cells by multiple mechanisms.

### Endosymbiosis with *R. pickettii* is required for immune evasion and virulence *in vivo*

Finally, we investigated the influence of bacterial-fungal symbiosis in Mucorales pathogenesis *in vivo*. Using our recently-established zebrafish (*Danio rerio*) model of infection (41), wild-type larvae were infected with either resting or swollen wild type or cured spores (**Supp. Figure 3A**). Over 96 hours post-infection (hpi), *R. microsporus* FP469-12 spores containing the endosymbiont killed a significant proportion of the larvae, with swollen spores being more virulent than resting (65% mortality at 96 hpi, compared to 40%, **Figure 7A**). In contrast, spores lacking the endosymbiont were completely avirulent, with neither swollen nor resting spores statistically different from mock injection in this model (*p*>0.05). This correlated with CFUs over time: While there were no differences in initial inocula, fungal CFUs from fish infected with endosymbiont-free spores were significantly reduced within just 2 hours and continued to be more rapidly cleared than wildtype spores over 96 hours **(Figure 7B**). This was independent of whether fish were infected with resting or pre-swollen spores, although in both cases, resting spores were more rapidly cleared (**Figure 7C**). Together, these data suggest that the endosymbiont aids immune evasion during the initial phase of infection, and that this is exacerbated by metabolic pre-activation of spores.

**Figure 7:**
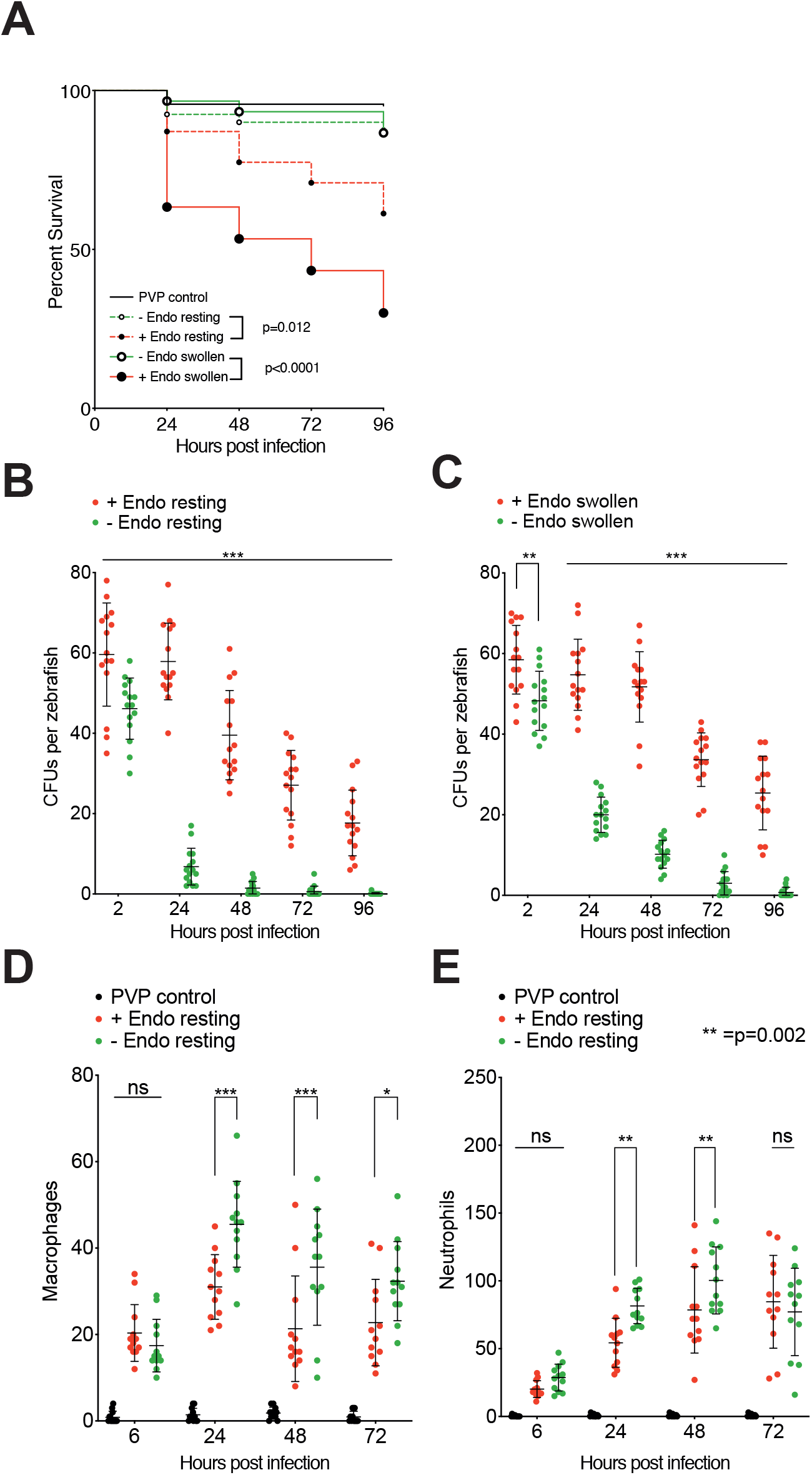
Effect of endosymbiont status on fungal infection of zebrafish. (A) Survival of AB wildtype zebrafish injected via the hindbrain with resting or swollen spores of *R. microsporus* FP469-12 in the presence or absence of endosymbiotic bacteria. PVP indicates mock-injected fish. Three biological replicates of populations of 10 fish each were examined (n=30). Statistical differences were determined using Mantel-Cox with Bonferroni’s correction for multiple comparisons (5% family-wise significance threshold = 0.025). (B) and (C) Effect of endosymbiont status on fungal survival (CFUs) following hindbrain injections of AB wild type zebrafish with (B) resting or (C) swollen *R. microsporus* FP469-12 spores. Three biological replicates of 5 fish per condition were examined (n=15). (D) and (E) show the effect of the endosymbiont on *in vivo* recruitment of macrophages and neutrophils to the site of infection. Supp. Figure 3 shows representative images and Supp. Figure 4 shows equivalent data with swollen spores. Statistical significance was assessed by Two-way ANOVA with Tukey’s correction for multiple comparisons or pairwise t-tests where sample number was unequal due to fish death, *=p<0.05; **=p<0.001; ***=p<0.0001 unless otherwise indicated.

Disease outcome correlates with the peak number of phagocytes at the site of infection (carrying capacity) in the first 24 hours, and successful infection control requires spore killing (17, 41). To investigate the role of the endosymbiont in modulating host defence, we quantified immune cell recruitment using transgenic zebrafish where either macrophages or neutrophils were fluorescently labelled (Tg(mpeg1:G/U:NfsB-mCherry or Tg(mpx:GFP)^i114^ respectively (66, 67)) (**Supp. Figure 3**). Zebrafish were therefore infected with wild-type or endosymbiont-free resting spores and the number of phagocytes recruited to the site of infection counted over the following 96 hours (for survival curves, see **Supp. Figure 4A and B**).

24 hours post-infection, both macrophage and neutrophil recruitment was significantly lower in larvae injected with spores containing the endosymbiont than those without (**Figure 7D and E**, p<0.0001). This difference was sustained over 48 hours, but became less significant at 72 hpi, most likely due to different disease progression as endosymbiont-free spores are cleared (**Figure 7A**). Similar results were observed with spores pre-swollen before injection (**Supp Figure 4C and D**). Whilst fish that failed to recruit macrophages after 24 hours did not survive until the end of the assay regardless of endosymbiont status, only macrophages recruited to endosymbiont-free infections efficiently contained the infection. Whilst it remains an open question whether recruitment is inhibited by secreted factors, or other changes in fungal biology mediated by the endosymbiont, the presence of *R. pickettii* enhances *R. microsporus* virulence *in vivo* by multiple mechanisms, leading to suppression of the immune response and fungal clearance.

As endosymbiont removal promoted spore clearance in zebrafish, we attempted to model the impact of antibiotic treatment immediately prior to or during infection on disease outcome. Concomitant treatment of the fish water with 60 μg/mL ciprofloxacin had no effect on fish survival or spore CFU upon infection with either resting or swollen spores (**Supp Figure 4E and F**). Unfortunately, it is not possible to differentiate whether this is due to lack of effect or insufficient drug permeability. Therefore, we instead performed a small proof of concept pilot study in immune-competent mice (n=5 per group). Mice were infected intra-tracheally with resting spores that had been untreated or treated with ciprofloxacin for 3 hours immediately prior to inoculation and CFUs measured after 4 and 48 hours. Similar to the rapid spore killing observed in fish (**Figure 7B**), within 4 hours, there was a reduction in CFUs recovered from mouse lungs infected with ciprofloxacin-treated resting spores (p=0.019) (**Figure 8A**). We ruled out a direct inhibitory effect of ciprofloxacin on fungal survival as swollen spores treated with ciprofloxacin were as resistant to host killing as untreated swollen spores (p=0.095) (**Figure 8C**). After 48 hours, whilst 3/5 mice infected with untreated resting spores remained positive by CFU, all 5 mice infected with ciprofloxacin-treated resting spores cleared the infection (**Figure 8B**). Whilst underpowered, this was statistically significant (p=0.0384, Chi-square test, Newcombe/Wilson with continuity correction, with 60% attributable risk). This effect was lost when mice were infected with pre-swollen spores, where any secreted factors are washed away prior to infection (**Figure 8C**). Therefore, whilst endosymbiosis most likely evolved to promote survival in the environment, it also confers protection from phagocytic immune cells in animals, demonstrating an important role in facilitating opportunistic infection.

**Figure 8:**
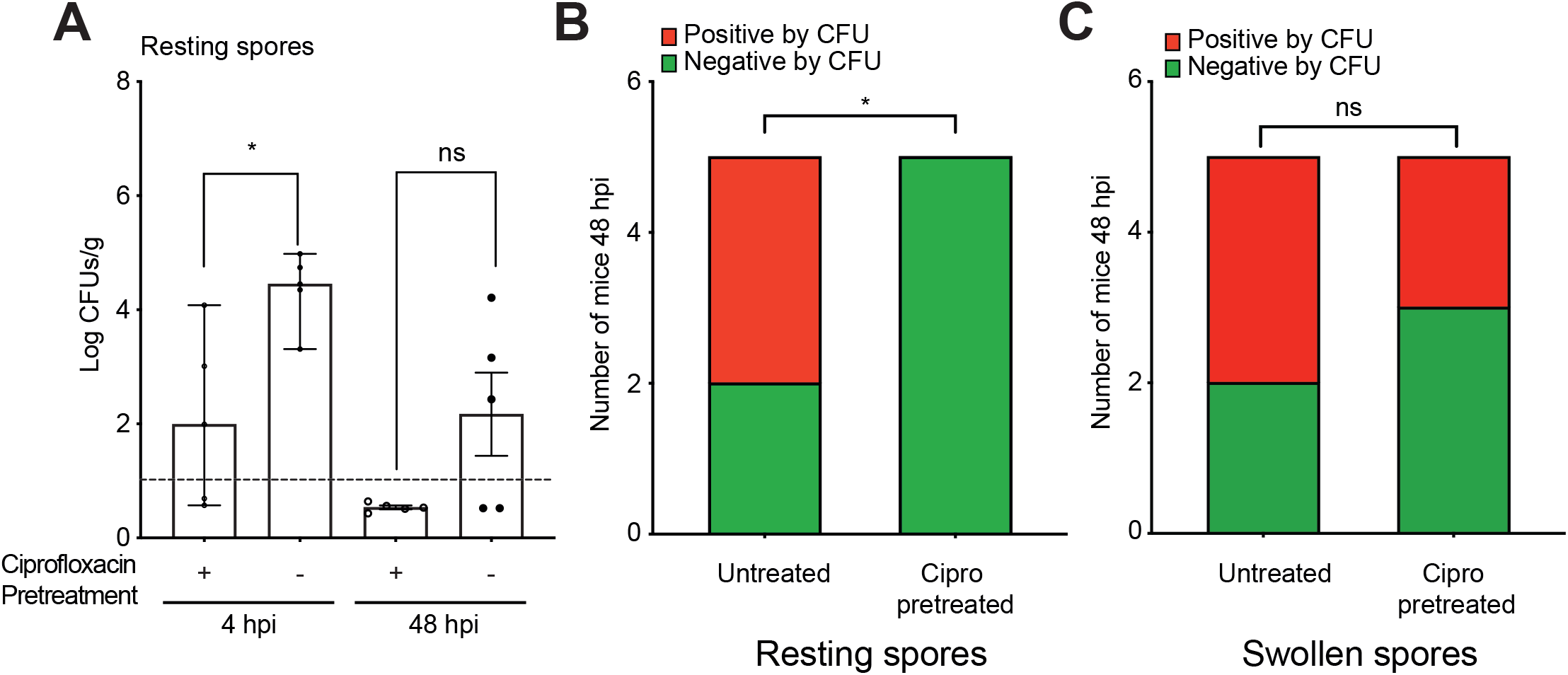
Ciprofloxacin treatment can modify *R. microsporus* FP469-12 infection in mice. (A) Fungal survival (CFU’s) at 4 and 48 following intra-tracheal infection of mice hours with resting spores. Where indicated, spores were pre-treated with 60 μg/ml Ciprofloxacin for 3hrs prior to infection. (n=5, significance determined by Mann-Whitney test). (B) and (C) Proportion of mice positive or negative (below the detection limit) for fungal CFU’s 48 hours post-infection. (B) shows data from infection with resting spores with and without ciprofloxacin pre-treatment, (C) is the same experiment performed with swollen spores. N=5 mice per condition, statistical significance assessed using a 2-sided Chi-squared test, and Attributable Risk was assessed using Newcombe/Wilson with continuity correction, *=p=0.384.

## Discussion

In this work, we describe an endosymbiosis between the opportunistic fungal pathogen *Rhizopus microsporus* and the gram-negative bacterium *Ralstonia pickettii*. Both are widely distributed in the environment, normally living in the soil where they will be preyed upon by microbial phagocytes, such as amoebae. A wide diversity of fungi harbour bacterial endosymbionts that influence fungal phenotypes related to metabolism, cell wall organization, development, and plant host colonization (35, 38, 39, 48, 64, 68–79). Endobacteria are also a source of potent mycotoxins that can influence fungal pathogenesis in plants and insects (39, 40, 48, 64, 80, 81). Here, we demonstrate a novel interaction, whereby endosymbiosis between *R. microsporus* and *R. pickettii* enables the biosynthesis and secretion of factors that inhibit both the growth of amoebae, and their ability to capture and kill other microbes. We propose the tri-kingdom interaction with amoebae provides an additional evolutionary driver for endosymbiosis, enabling both bacteria and fungus to evade predation in the environment.

Consistent with long-term mutualism, the presence of the *R. pickettii* endosymbiont caused widespread changes to its host; endosymbiont removal altered *R. microsporus* cell wall structure and reduced resistance to both reactive nitrogen and oxygen species. Mondo *et al.*, showed that bacterial endosymbionts can impact fungal sporulation and development through controlling *ras2*, a key GTPase that regulates fungal sexual reproduction (38). Consistent with this, we also observed reduced sporulation and overall fitness after repeated passage of endosymbiont-free fungi on SDA plates with ciprofloxacin (data not shown). Loss of the endosymbiont also increased sensitivity to the front-line antifungal treatment Amphotercin B. This indicates altered ergosterol content, and suggests an additional way in which the endosymbiont may influence the outcome of human infections.

In our companion work examining the transcriptional response of *R. microsporus* to endosymbiosis with *R. pickettii*, we observed limited changes during growth in DMEM (82). Loss of the endosymbiont however profoundly changed the response of *R. microsporus* to phagocytosis, causing altered regulation of 359 genes, including inappropriate repression of genes involved in siderophore activity and iron scavenging. We observed a similar global change in the intensity of the fungal response to phagocytosis in a cured isolate of *R. delemar* lacking an unculturable endosymbiont with high 16S sequence homology to other unculturable bacteria (Accession number MK573034)(82). This provides additional evidence for a general role for phagocyte interactions in driving fungal/bacterial endosymbiosis.

Most studies of *R. microsporus* secreted mycotoxins have focussed on the rhizoxin and rhizonin molecules generated by *B. rhizoxina* endosymbionts (45, 48, 64, 74). Although *Ralstonia* and *Burholderia* spp are related and share the same niche, the anti-amoebic activity of the *R. microsporus* FP469-12 endosymbiosis differs in several important ways. Firstly, whilst *B. rhizoxina* alone is sufficient for rhizoxin biosynthesis (40, 64), media conditioned with the isolated *R. pickettii* endosymbiont from *R. microsporus* FP469-12 was ineffective against either *D. discoideum* or macrophages, indicating in this case both fungi and bacteria are necessary for the secreted bioactivity.

The mechanism of action also differs; whilst rhizoxin potently depolymerises the microtubule cytoskeleton, this appeared unaffected in amoebae treated with *R. microsporus* FP469-12-condiditioned medium. Interestingly, whilst rhizoxin is effective at picomolar concentrations in plants and mammalian cells (47, 64), *D. discoideum* were several orders of magnitude more resistant to Rhizoxin D. This is most likely due to the highly active multi-drug efflux pumps in amoebae that presumably evolved for precisely this reason – to protect against environmental toxins. Whilst we cannot exclude more subtle effects on, for example, microtubule motors, almost complete depolymerisation of the microtubule array was required to inhibit growth to a similar extent as conditioned medium. Future studies are needed to identify the active molecule and target, but it seems likely that amoebae growth and phagosome maturation are perturbed by a novel mechanism.

An interesting additional aspect to the relationship between environmental fungi, bacteria and amoebae is that bacteria can also form endosymbiotic relationships with amoebae (83, 84). In particular several *Burkholderia* spp have been identified as stable symbionts associated with environmental isolates *D. discoideum* (85–87). The relationships between amoeba and symbionts are complex and not always obvious, with both costs and benefits to both organisms described (84, 87, 88). Nonetheless, the different symbiotic relationships between bacteria, fungi and amoebae, within the same environmental niche, underlines the multifactorial interactions occurring in complex microbial communities.

As the majority of fungi live in the environment and do not normally infect animals, how they evolved the mechanisms that allow them to evade the human immune system and cause opportunistic infections is unclear. As phagocytosis and phagosome maturation are highly conserved across evolution (89), it has been proposed that opportunistic fungal virulence derives from environmental interactions with amoebae (53, 90). Our data show that the presence of the endosymbiont confers resistance to macrophages in a similar manner to amoebae, inhibiting both phagocytosis and killing. The presence of the endosymbiont is also essential for virulence in a zebrafish model of infection. Whilst the widespread changes in fungal physiology that occur upon endosymbiont removal indicate that evading amoebae is not the sole driver of endosymbiosis, we suggest it is an important contributary factor, and significant for the evolution of virulence.

Whilst a broadening diversity of basal saprophytic fungi are a major emerging cause of clinically important superficial and disseminated fungal infections, the role of endosymbionts in pathogenesis has yet to be clinically defined (10, 91). Two groups recently reported that although bacterial endosymbionts, including rhizoxin and rhizonin producers, are widely found among mucormycete clinical isolates, these endosymbionts did not impact pathogenesis during mucormycosis in both mouse and fly models (35, 45). In a diabetic mouse model infected with *Rhizopus* isolates harbouring rhizoxin-producing *Burkholderia* endosymbionts, no impact of treatment with ciprofloxacin 24 hours post-infection was observed (35). In contrast, we found pretreatment of resting spores with ciprofloxacin before infecting immune-competent mice significantly reduced CFUs retrieved after 4 hours and spores could no longer be recovered after 48 hours. This benefit was however lost when spores were pre-swollen prior to infection, indicating that to be therapeutically effective, endosymbiont removal would need to occur early in the infection. Whilst our study only used a small number of mice (5 per group) and is therefore prone to the challenges associated with underpowered models of infection, similar patterns were observed in zebrafish, with much larger sample sizes. Whilst most studies to date have focussed on *Burkholderia* spp endosymbionts and the rhizoxin and rhizonin mycotoxins, our work indicates a broader diversity of both endosymbionts and virulence mechanisms. The relevance of endosymbiosis to human infections therefore needs readdressing, using larger-scale studies of the prevalence and diversity of endosymbionts in both clinical and environmental isolates.

Our findings highlight the potential importance of bacterial symbiosis in *R. microsporus* pathogenesis. However, we urge caution in extending our findings to all Mucorales or fungal endosymbionts. While a number of studies have demonstrated general features common to host-Mucorales interactions, there are also likely to be species-specific aspects. In addition, not all clinical isolates profiled harbour endosymbionts, suggesting they may augment, but are not constitutively required for, pathogenesis (35). Overall however, this work points to a new role for bacterial endosymbionts in fungal immune evasion, driven by environmental interactions with amoebae. The observation that endosymbiont elimination impacts phagocytosis, cell wall organization, stress resistance and antifungal drug susceptibility as well as resistance to macrophage-mediated killing raises the prospect that endosymbiont-targeted treatments may be useful in the treatment and management of a subset of opportunistic fungal infections.

## Methods

### For full detailed methods see the Supplementary Information

#### Data and material availability

Strains unique to this study will be made available to other users upon request. Genomic and transcriptomic sequencing data is available through the Sequence Read Archive (SRA) on NCBI and accession codes will be provided and made public upon publication.

#### *R. microsporus* strain and culture

*Rhizopus microsporus* strain FP469-12.6652333 was isolated from a human infection at the Queen Elizabeth Hospital Birmingham Trauma Centre by Dr. Deborah Mortiboy. This was routinely cultured on Sabouraud dextrose agar (SDA) (EMD Millipore co-operation) or potato dextrose agar (PDA) to induce sporulation at room temperature. Routine liquid cultivation was performed in 500 mL conical flasks containing 250 mL of serum free Dulbecco’s modified eagle’s media (with 1% penicillin/streptomycin and L-glutamine) (DMEM). Fungi grown in this way was used for metabolic activation of spores, to prepare swollen spore supernatants, and for DNA isolation.

Conditioned medium was generated by incubating at 4×10^8^ spores/mL in 250 mL of either serum free DMEM (for macrophage experiments) or HL5 medium (Formedium, for *D. discoideum* experiments) at 37°C with shaking at 200 rpm for 4 h. The medium was then centrifuged at 3990 rpm for 5 mins and the supernatant filter sterilised through a 0.45 μm filter.

#### Macrophage cell line culture

J774A.1 murine macrophage-like cells were maintained in DMEM supplemented with 10% foetal bovine serum, 1% Streptomycin (100 μg/mL), penicillin (100 U/mL), and 1% L-glutamine (2 mM). The cells were cultivated in a humidified environment at 37°C enriched with 5% CO_2_ and used between 3 and 15 passages after thawing.

#### *D. discoideum* culture

All experiments were performed using the Ax2 axenic strain, routinely subcultured in HL5 medium (Formedium) at 22 °C. For growth curves, cells were seeding cells at 1 × 10^5^/ml in the appropriate medium and counted at regular intervals until out of exponential growth. GFP-α tubulin was expressed using extrachromosomal vector pJSK336 (92); cells were transformed by electroporation and transformants selected in 20 μg/mL G418. Using ImageJ (93), maximum intensity projections of images from a Perkin-Elmer Ultraview VoX (60X 1.4NA objective) were then used to manually draw around the tips of the microtubule array, and calculate the areas covered as a proportion of the whole cytosol.

#### Spore viability

Resistance to killing by macrophages was assessed as following phagocytosis at the indicated time points. Macrophages were lysed with 1 mL sterile water and aggressively washed to collect adherent spore cells. The lysate was serially diluted and 5 *μ*L plated out on SDA for CFUs.

### Virulence assays

#### Zebrafish maintenance and infection

Infections were performed in wild type AB zebrafish, as well as GFP-neutrophil (Tg(mpx:GFP)^i114^) lines (66). Macrophage-specific mCherry expression was achieved by crossing Tg(mpeg1:Gal4-FF)^gl25^ with Tg(UAS-E1b:NfsB.mCherry)^c264^, referred to as Tg(mpeg1:G/U:NfsB-mCherry) (67). Zebrafish were cultivated under a 14 h-10 h light-dark cycle at 28°C at the University of Birmingham (BMS) Zebrafish Facility. All zebrafish care protocols and experiments were performed in accordance with the UK animals (scientific procedures) act, 1986. Following collection of the eggs, at 40-60 eggs per 25 mL E3 media (plus 0.1% methylene blue and 1 mg/mL 1-phenyl- 2- thiourea (PTU)) for 24 h and larvae were incubated at 28°C. PTU at this concentration prevented melanization of the embryos. All media and reagents used was purchased from Sigma Aldrich unless otherwise mentioned.

Prior to injection, spores collected from 10 day old cultures on SabDex plates were washed 3x in PBS and stained with Calcofluor White in 0.1 M of NaHCO_3_ for 30 min. Swollen spores were pre-incubated in DMEM at 37°C, 200 rpm for 4 hr prior to staining. Spores were the washed 3 times in PBS, and resuspended at 10^8^ spores/mL in 10% (w/v) polyvinylpyrrolidone-40 (PVP) in PBS with 0.05% phenol red in ddH_2_O. PVP was used as a carrier medium to increase density and prevent clogging of microinjection needles. Zebrafish were injected in accordance to a protocol by Brothers *et al*., and zebra fish development assessment in accordance to Kimmel *et al*. (94, 95). The fish were injected at prim-25 stage following manual dechorionation and anesthesia with 160 μg/mL of Tricaine in ddH_2_O.

Micro-injection was performed with 2 nL of 10% (PVP) in PBS or *R. microsporus* spore suspension through the otic vesicle into the hind brain to achieve an inoculum dose of approximately 50-100 spores per fish. Following injection, larvae were anesthetized with 160 μg/mL Tricaine in E3 media in a 96 well plate and screened by fluorescence microscopy, (Zeiss Axioserver Zi microscope equipped with Aptome system) for the presence of spores. Only larvae with approximate correct inoculum were selected, and transferred to individual wells of a 24 well plate containing E3 media (plus 0.1% methylene blue ± 60 μg/mL Ciprofloxacin). The fish were monitored over a period of 96 hours post infection for survival whereupon they were killed by 1600 μg/mL Tricaine overdose and treated with bleach overnight before disposal.

#### Recovery of fungal spores

Following injection, 5 fish per biological repeat were euthanized with 1600 μg/mL Tricaine at each timepoint, before homogenisation in 100 μL of E3 media containing penicillin-streptomycin (5000 U/mL-5 mg/mL) and gentamicin (10 mg/mL) using pellet pestles. Extracts were then plated out on SDA containing 100 U/mL-100 μg/mL penicillin-streptomycin and 30 μg/mL gentamicin, incubated at room temperature for between 24 and 48 h and colony forming units (CFUs) examined.

#### Phagocyte recruitment

To quantify phagocyte recruitment into the hindbrain, Tg(mpx: GFP)^i114^ or Tg(mpeg1:G/U:NfsB-mCherry) transgenic zebrafish (66, 67) were injected as above. At each timepoint, fish were imaged using an Axio Observer Z1 microscope equipped with Apotome (Carl Zeiss), to reconstruct the 3D volumes. Positive phagocyte recruitment was defined by accumulation of >10 neutrophils or macrophages to the site of infection. At least 3 biological repeats were performed with 5 fish per condition to give a total of 15 fish per group.

#### Mouse model

Immunocompetent CD-1 20-23g male mice (n=12 per group) were randomly assigned to one of two treatment groups and the groups housed in IVCs. *Rhizopus microsporus* spores (FP469-12.6652333) were collected from Sabouraud agar plates in 10 mL of PBS + 0.01% Tween 20, washed once with endotoxin-free PBS and resuspended in serum-free DMEM at 4×10^7^/ml. To mimic the early stages of infection, swollen spores were pre-germinated in serum-free DMEM with or without Ciprofloxicin (60 μg/mL) for up to 3 hrs at 37°C, 200 rpm, sufficient to swell but not form germ tubes. Mice were infected intratracheally with 25μl serum-free DMEM containing 10^6^ resting or swollen spores. Mice (n=5 per group) were sacrificed by pentobarbital overdose and lungs were collected at two different time points: 4 h or 48 h post infection. Lungs were homogenization in 2 ml of PBS, and 200 μl were plated directly from the concentrated samples and also from serial dilutions onto potato dextrose agar + 0.1% Triton plates and incubated at 37 °C. Innoculum was verified via lung CFU from two mice directly after infection. Data were tested for normality using Shapiro-Wilk and analysed using the Mann-Whitney U test. Data are reported as Median with 95% CI.

#### Ethical statement

All maintenance protocols and experiments were performed in accordance to Animals (scientific procedures) Act, 1986 as required by United Kingdom (UK) and European Union (EU). All work was performed under appropriate Biosafety Level 2 conditions (BSL2). All zebrafish care and experimental procedures were conducted according to Home Office legislation and the Animals (Scientific Procedures) Act 1986 (ASPA) under the Home Office project license 40/3681 and personal licenses l13220C2C to Kerstin Voelz and lCDB92D64 to Herbert Itabangi. Mouse studies were approved by the Institutional Animal Care and Use Committee of the Los Angeles Biomedical Research Institute at Harbor-UCLA Medical Center, according to the NIH guidelines for animal housing and care under protocol 11671.

## Acknowledgements

We are grateful to Dr. Deborah Mortiboy of the Queen Elizabeth Hospital Birmingham Trauma Centre for isolation and initial identification of *Rhizopus microsporus* FP469-12.6652333, to Zoe Reading and Bradley Pollard for their contributions to the work during their research projects in the Voelz lab, and to the University of Aberdeen Microscopy and Histology Facility.

## Funding support

This work was supported by a Wellcome Trust Seed award to KV (108387/Z/15/Z). HI is supported by the Wellcome Trust Strategic Award in Medical Mycology and Fungal Immunology (097377). PSC is supported by a BBSRC MIBTP PhD Studentship (BB/M01116X/1). XZ is supported by a Studentship from the Darwin Trust of Edinburgh. PJM is supported by the UK Biotechnology and Biological Research Council (BB/S010122/1)

ASI is supported by Public Health Service grants from the National Institutes of Allergy and Immunology R01 AI063503. GDB is funded by Wellcome Trust (102705) and MRC Centre for medical mycology (MR/N006364/2).ERB was supported by the UK Biotechnology and Biological Research Council (BB/M014525/1) and a Sir Henry Dale Fellowship jointly funded by the Wellcome Trust and the Royal Society (211241/Z/18/Z). JSK is funded by Royal Society University Research Fellowship UF140624. GPS is funded by Royal Society grant RG150439.

## Author Contributions

HI designed the experiments, performed the work, performed the analysis, and wrote the manuscript.

PSC designed the RNASeq experiments, collected the data, performed the analysis, contributed to the interpretation, and contributed to the manuscript.

XZ contributed to the acquisition and analysis of data and contributed to the manuscript.

BP contributed to the acquisition and analysis of data and contributed to the manuscript.

II contributed to the acquisition and analysis of data and contributed to the manuscript.

MP contributed to the acquisition and analysis of data and contributed to the manuscript.

ZR contributed to the acquisition and analysis of data and contributed to the manuscript.

JC contributed to the acquisition and analysis of data and contributed to the manuscript.

PJM contributed to the experimental design and acquisition and analysis of data and contributed to the manuscript.

TG contributed to the acquisition and analysis of data and contributed to the manuscript.

YG contributed to the acquisition and analysis of data and contributed to the manuscript.

LL contributed to the acquisition and analysis of data and contributed to the manuscript.

BP contributed to the acquisition and analysis of data and contributed to the manuscript.

ASI contributed to the experimental design and interpretation and contributed to the manuscript.

GB contributed to the experimental design and interpretation and contributed to the manuscript.

FFT contributed to the experimental design and interpretation and contributed to the manuscript.

JK contributed to the experimental design, analysis and interpretation of data and wrote the manuscript.

ERB contributed to the analysis and interpretation of data and wrote the manuscript.

KV conceived and designed the experiments, contributed to the analysis and interpretation of data, and wrote the manuscript.

## Itabangi et al. Supplementary methods

### *R. microsporus* growth and manipulation

Endosymbionts were isolated by submerging a small mycelial pellet from a 2-4 day old liquid culture into 500 *μ*L of 2000 units/mL Lyticase enzyme (Sigma-Aldrich) in distilled water and incubated at 25°C for 2.5 h. The mycelia were further disrupted by pipetting and vortexing before gradient centrifugation at 13200 rpm for 30 mins. 10 *μ*L of supernatant was then aseptically plated on nutrient agar and incubated at 30°C until bacterial or fungal growth was observed. Bacterial colonies were then sub cultured in LB broth.

Endosymbiont-free (cured) fungal strains were obtained by cultivating the fungi in the presence of 60 μg/mL ciprofloxacin for a month. Fungi were then maintained on ciprofloxacin plates for 3 months before the endosymbiont was entirely cleared. Absence of the bacteria was routinely confirmed through SYTO9 staining, and PCR screening for bacterial 16S rDNA, as described below.

Spore stress resistance was measured by incubating at 10^5^ spores/mL in serum-free DMEM containing the indicated concentrations of hydrogen peroxide (H_2_O_2_), sodium nitrite (NaNO_2_, Fisher scientific), sodium dodecyl sulphate (SDS) (Fisher scientific), sodium chloride (NaCl, Sigma-Aldrich) or Amphotericin B (AmB, Sigma Aldrich). The spores were incubated at 37oC and 5% CO_2_ for 24 or 48h before CFUs were determined by plating serial dilutions on SDA agar.

### Cell wall analysis

For microscopy, 2×10^7^ spores/ml were incubated for 30 minutes at room temperature in PBS containing either 100 μg/ml fluorescein isothiocynate isomer 1 (FITC) (Sigma-Aldrich) in 0.1 M sodium bicarbonate buffer (pH 7.45) (Sigma Aldrich), 25-50 μg/mL RhTRITC-concanavalin A (ThermoFisher scientific) or 250 μg/ml calcofluor white (Sigma Aldrich). Spores were then imaged at 63x magnification on a Zeiss Axio Observer Z1 equipped with structured illumination (Apotome) using a Flash 4 sCMOS camera (Hamamatsu) and the same exposure across samples. Staining intensity was quantified for each cell using ImageJ to measure relative fluorescence within computationally identified regions of interest.

For flow cytometry, resting *R. microsporus* FP 469-12 spores were pre-germinated for the indicated times in serum-free DMEM at 37°C, 200 rpm, before fixation in 4% methanol-free formaldehyde, and incubation with either 0.01 μg/ml Dectin-1 IgG antibody, washed, and 1:200 goat anti-mouse IgG-488 secondary antibody (β-glucan); Calcofluor white (CFW, chitin; 250 μg/ml); or ConcanavalinA (ConA-488; 50 μg/ml). At least 10,000 cells were measured using an Attune NxT flow cytometer and compared to a pooled secondary-only control. Data are representative of three independent experiments.

### TEM

*R. microsporus spores* were collected and allowed to swell as described above. Samples were processed via high-pressure freezing using a Bal-Tec HPM 010 high-pressure freezer (Boeckler Instruments, Tucson, AZ) and transferred to an RMC FS-7500 freeze substitution unit (Boeckler Instruments, Tucson, AZ) before substitution in 2% osmium tetroxide, 1% uranyl acetate, 1% methanol, and 5% water in acetone. Samples were then transitioned from −90°C to room temperature over 2 to 3 days, rinsed in acetone, and embedded in LX112 epoxy resin (Ladd Inc., Burlington, VT). Ultrathin sections of 70 to 80 nm were cut on a Leica Ultracut UC7 microtome, stained with uranyl acetate followed by lead citrate, and viewed on a JEOL 1200EX transmission electron microscope at 80 kV.

### SYTO9 staining of bacterial endosymbionts

A small sample of mycelia pellet was aseptically submerged into 200 mL of 0.85% sodium chloride (NaCl) for 1 h, washed 2x in PBS and stained with the Live/Dead BacLight bacterial viability kit (Thermofisher Scientific). Mycelia were incubated in stain solution for 15 minutes, fixed with 4% PFA, mounted on glass slides and imaged using a 63x oil objective under phase contrast on a Zeiss Axio Observer Z1 microscope.

### Screening and identification of bacterial endosymbiont

PCR screening for the presence of endosymbionts was performed using Universal primers (35, 40) to amplify the 1.5 kb band for 16S rDNA from genomic DNA isolated from mucormycetes (5’CCGAATTCGTCGACAACAGAGTTTGATCCTGGCTCAG3’/ 5’CCCGGGATCCAAGCTTACGGCTACCTTGTTACGACTT 3). DNA was obtained from 10^7^ spores in a screw cup with beads and homogenised using a bead beater (Bertin technologies) at 6500x *g* for 1 min and DNA extracted with a DNeasy® powerlyzer® microbial kit (Qiagen) following the manufacturer’s instructions. PCR was performed using Phusion kit (New England Biolabs) and 35 cycles, of 95° for 2 min, 60°C for 30s, and 72°C for 1 min. The resulting PCR product was then sequenced, identifying. *R. pickettii*. For full-genome sequencing, bacterial genomic DNA was extracted using the DNeasy PowerLyzer Microbial Kit (Qaigen) and sequenced by MicrobesNG (University of Birmingham, UK). The sample was identified by kraken (v1) as being highly similar to *R. picketti* J12. Whole genome sequence data is publicly available (see data availability statement).

### Phagocytosis assays

For phagocytosis by macrophages, 1 ×10^5^ J774A.1 cells (94) were seeded per well a 24 well plate in DMEM and incubated overnight. Next day, cells were washed 2x in pre-warmed PBS, before incubation with 1 mL pre-warmed serum-free DMEM at 37°C +5% CO_2_ for 1 h. Cells were then washed 2x with PBS, and FITC pre-stained spores at 5×10^5^ spores/mL in sfDMEM added to give a multiplicity of infection (MOI) of 1:5. The co-culture was incubated 37°C +5% CO_2_ for 1 h before fixation in 4% PFA for 15 min, and washing with PBS prior to counter staining with Concanavalin A (Con-A) or calcoflour white (CFW) for 30 min. Cells were then washed in PBS and imaged on a Nikon T1 microscope. Uptake rate was quantified as the number of phagocytes containing at least one spore, >1000 phagocytes assessed per replicate.

Phagocytosis of UV killed spores was performed as above, except prior to addition, spores were irradiating twice for 15 min in a UV PCL-crosslinker at 1200 *μ*J/cm^2^ in PBS, cooling on ice between treatments as previously described (17). Successful killing was confirmed by plating for CFUs. Latex beads, *C. albicans* SC5314 and *S. cerevisisae* AM13/001 were all processed similarly to spores including washing with PBS prior to addition to the macrophages.

Phagocytosis by *D. discoideum* was measured by incubating 2 × 10^6^ amoebae in 2ml untreated or conditioned HL5 in 3cm glass-bottomed microscopy dishes for 1 hour, prior to addition of 1 × 10^7^ TRITC-labelled heat killed *S. cerevisiae* (prepared as in (95)). After 30 minutes, fluorescence of unengulfed yeast was quenched by addition of 100 μl 0.4% trypan blue solution (Sigma Aldrich), before fluorescence microscopy and quantification of number of yeast engulfed per amoebae, counting >100 cells quantified per sample.

**Phagosomal proteolysis** was measured as previously described using 3 μm silica beads co-labelled with DQ-green BSA and Alexa594 (performed as in (96)). Amoebae were washed twice in LoFlo medium (Formedium), before resuspension at 3 × 10^6^/ml in HL5 medium and 100 μl/ well seeded in clear-bottomed black-walled 96-well plates (Greiner). After 2 hours, 10 μl reporter beads were added to triplicate wells at a bead:cell ratio of 1:2, and the plate spun down at 1,200 rpm for 10 seconds to synchronise uptake. Free beads were then removed by tapping the inverted plate on a paper towel and washing twice in HL5. 100 μl conditioned, or control media was then added to wells before the plate was placed in a plate reader, and fluorescence measured at 500/520nm and 594/630nm (excitation/emission) every minute. For pre-treatment with conditioned medium, it was added 30 minutes prior to bead addition, and used in all subsequent washes/incubations. Proteolysis was calculated by the increase in DQ-BSA fluorescence over time, normalised to bead uptake, determined by Alexa594 fluorescence.

**Phagosomal killing** was performed as in (97, 98) following the quenching of GFP-expressing *Klebsiella pneumoniae* fluorescence (gift from Pierre Cosson, University of Geneva). 10 μl of a saturated overnight bacterial culture in LB was diluted in 280 μl HL5 (or conditioned medium) and allowed to settle as a drop in a glass-bottomed microscopy dish (Mat-Tek). After 15 minutes, 1.5 ml of *D. discoideum* culture at 1 × 10^6^ cells/ml was carefully added and GFP and bright-field imaged recorded every 20 seconds for 40 minutes at 20x magnification. Movies were then manually analysed to determine the time of GFP-quenching post-engulfment.

### Statistics

All data was analysed in Graphpad Prism 7 using the nonparametric, Mann-Whitney U tests, and log rank tests, with Dunn’s correction for multiple comparisons where appropriate and as indicated in the figure legends and main text. Differences with *p* value ≤0.05 were considered significant. All experiments were performed in triplicate, with the number of events measured indicated in figure legends. Data are plotted as Box-and× Whiskers with maximum and minimum values, except where data shown are percentages or relative values.

## Notes

### Competing Interest Statement

The authors have declared no competing interest.

### Summary of Updates

Revised to reflect reviewer feedback

## References

1. Erwig LP & Gow NA (2016) Interactions of fungal pathogens with phagocytes. Nat Rev Microbiol 14(3):163–176.

2. Novohradska S, Ferling I, & Hillmann F (2017) Exploring Virulence Determinants of Filamentous Fungal Pathogens through Interactions with Soil Amoebae. Frontiers in cellular and infection microbiology 7:497.

3. Chamilos G, et al. (2010) Generation of IL-23 producing dendritic cells (DCs) by airborne fungi regulates fungal pathogenicity via the induction of T(H)-17 responses. PLoS One 5(9):e12955.

4. Petraitis V, et al. (2013) Increased virulence of Cunninghamella bertholletiae in experimental pulmonary mucormycosis: correlation with circulating molecular biomarkers, sporangiospore germination and hyphal metabolism. Med Mycol 51(1):72–82.

5. Roden MM, et al. (2005) Epidemiology and outcome of zygomycosis: a review of 929 reported cases. Clinical infectious diseases: an official publication of the Infectious Diseases Society of America 41(5):634–653.

6. de Hoog S, Ibrahim AS, & Voigt K (2014) Zygomycetes: an emerging problem in the clinical laboratory. Mycoses 57 Suppl 3:1.

7. Douglas AP, Chen SC, & Slavin MA (2016) Emerging infections caused by non-Aspergillus filamentous fungi. Clinical microbiology and infection: the official publication of the European Society of Clinical Microbiology and Infectious Diseases 22(8):670–680.

8. Gomes MZ, Lewis RE, & Kontoyiannis DP (2011) Mucormycosis caused by unusual mucormycetes, non-Rhizopus, -Mucor, and -Lichtheimia species. Clin Microbiol Rev 24(2):411–445.

9. Kontoyiannis DP & Lewis RE (2006) Invasive zygomycosis: update on pathogenesis, clinical manifestations, and management. Infectious disease clinics of North America 20(3):581–607, vi.

10. Ibrahim AS & Voelz K (2017) The mucormycete-host interface. Current opinion in microbiology 40:40–45.

11. Skiada A, Rigopoulos D, Larios G, Petrikkos G, & Katsambas A (2012) Global epidemiology of cutaneous zygomycosis. Clin Dermatol 30(6):628–632.

12. Blyth CC, et al. (2014) Consensus guidelines for the treatment of invasive mould infections in haematological malignancy and haemopoietic stem cell transplantation, 2014. Internal medicine journal 44(12b):1333–1349.

13. Medwid RD & Grant DW (1984) Germination of Rhizopus oligosporus Sporangiospores. Appl Environ Microbiol 48(6):1067–1071.

14. Turgeman T, et al. (2016) The Role of Aquaporins in pH-Dependent Germination of Rhizopus delemar Spores. PLoS One 11(3):e0150543.

15. Thau N, et al. (1994) rodletless mutants of Aspergillus fumigatus. Infect Immun 62(10):4380–4388.

16. Carrion SdJ, et al. (2013) The RodA hydrophobin on Aspergillus fumigatus spores masks dectin-1- and dectin-2-dependent responses and enhances fungal survival in vivo. Journal of Immunology (Baltimore, Md: 1950), 191(5):2581–2588.

17. Voelz K, Gratacap RL, & Wheeler RT (2015) A zebrafish larval model reveals early tissue-specific innate immune responses to Mucor circinelloides. Dis Model Mech 8(11):1375–1388.

18. Waldorf AR & Diamond RD (1985) Neutrophil chemotactic responses induced by fresh and swollen Rhizopus oryzae spores and Aspergillus fumigatus conidia. Infect Immun 48(2):458–463.

19. Chamilos G, et al. (2008) Drosophila melanogaster as a model host to dissect the immunopathogenesis of zygomycosis. Proc Natl Acad Sci U S A 105(27):9367–9372.

20. Andrianaki AM, et al. (2018) Iron restriction inside macrophages regulates pulmonary host defense against Rhizopus species. Nat Commun 9(1):3333.

21. Li CH, et al. (2011) Sporangiospore size dimorphism is linked to virulence of Mucor circinelloides. PLoS Pathog 7(6):e1002086.

22. Jennessen J, Schnurer J, Olsson J, Samson RA, & Dijksterhuis J (2008) Morphological characteristics of sporangiospores of the tempe fungus Rhizopus oligosporus differentiate it from other taxa of the R. microsporus group. Mycol Res 112(Pt 5):547–563.

23. Rappleye CA, Eissenberg LG, & Goldman WE (2007) Histoplasma capsulatum alpha-(1,3)-glucan blocks innate immune recognition by the beta-glucan receptor. Proc Natl Acad Sci U S A 104(4):1366–1370.

24. Garfoot AL, Shen Q, Wuthrich M, Klein BS, & Rappleye CA (2016) The Eng1 beta-Glucanase Enhances Histoplasma Virulence by Reducing beta-Glucan Exposure. MBio 7(2):e01388–01315.

25. Ballou ER, et al. (2016) Lactate signalling regulates fungal beta-glucan masking and immune evasion. Nat Microbiol 2:16238.

26. O’Meara TR & Alspaugh JA (2012) The Cryptococcus neoformans capsule: a sword and a shield. Clin Microbiol Rev 25(3):387–408.

27. Aimanianda V & Latge JP (2010) Fungal hydrophobins form a sheath preventing immune recognition of airborne conidia. Virulence 1(3):185–187.

28. Aimanianda V, et al. (2009) Surface hydrophobin prevents immune recognition of airborne fungal spores. Nature 460(7259):1117–1121.

29. Dagenais TR, et al. (2010) Aspergillus fumigatus LaeA-mediated phagocytosis is associated with a decreased hydrophobin layer. Infect Immun 78(2):823–829.

30. Chrisman CJ, Alvarez M, & Casadevall A (2010) Phagocytosis of Cryptococcus neoformans by, and nonlytic exocytosis from, Acanthamoeba castellanii. Applied and environmental microbiology 76(18):6056–6062.

31. Lecointe K, Cornu M, Leroy J, Coulon P, & Sendid B (2019) Polysaccharides Cell Wall Architecture of Mucorales. Front Microbiol 10:469.

32. Antachopoulos C, Demchok JP, Roilides E, & Walsh TJ (2010) Fungal biomass is a key factor affecting polymorphonuclear leucocyte-induced hyphal damage of filamentous fungi. Mycoses 53(4):321–328.

33. Schmidt S, et al. (2013) Rhizopus oryzae hyphae are damaged by human natural killer (NK) cells, but suppress NK cell mediated immunity. Immunobiology 218(7):939–944.

34. Schmidt S, Schneider A, Demir A, Lass-Florl C, & Lehrnbecher T (2016) Natural killer cell-mediated damage of clinical isolates of mucormycetes. Mycoses 59(1):34–38.

35. Ibrahim AS, et al. (2008) Bacterial endosymbiosis is widely present among zygomycetes but does not contribute to the pathogenesis of mucormycosis. The Journal of infectious diseases 198(7):1083–1090.

36. Kobayashi DY & Crouch JA (2009) Bacterial/Fungal interactions: from pathogens to mutualistic endosymbionts. Annu Rev Phytopathol 47:63–82.

37. Spraker JE, Sanchez LM, Lowe TM, Dorrestein PC, & Keller NP (2016) Ralstonia solanacearum lipopeptide induces chlamydospore development in fungi and facilitates bacterial entry into fungal tissues. ISME J 10(9):2317–2330.

38. Mondo SJ, et al. (2017) Bacterial endosymbionts influence host sexuality and reveal reproductive genes of early divergent fungi. Nat Commun 8(1):1843.

39. Partida-Martinez LP & Hertweck C (2005) Pathogenic fungus harbours endosymbiotic bacteria for toxin production. Nature 437(7060):884–888.

40. Partida-Martinez LP, et al. (2007) Rhizonin, the first mycotoxin isolated from the zygomycota, is not a fungal metabolite but is produced by bacterial endosymbionts. Appl Environ Microbiol 73(3):793–797.

41. Inglesfield S, et al. (2018) Robust Phagocyte Recruitment Controls the Opportunistic Fungal Pathogen Mucor circinelloides in Innate Granulomas In Vivo. MBio 9(2).

42. Gebremariam T, et al. (2014) CotH3 mediates fungal invasion of host cells during mucormycosis. J Clin Invest 124(1):237–250.

43. Baldin C & Ibrahim AS (2017) Molecular mechanisms of mucormycosis-The bitter and the sweet. PLoS Pathog 13(8):e1006408.

44. Tominaga Y & Tsujisaka Y (1981) Investigation of the Structure of Rhizopus Cell Wall with Lytic Enzymes. Agricultural and Biological Chemistry 45(7):1569–1575.

45. Partida-Martinez LP, Bandemer S, Ruchel R, Dannaoui E, & Hertweck C (2008) Lack of evidence of endosymbiotic toxin-producing bacteria in clinical Rhizopus isolates. Mycoses 51(3):266–269.

46. Prota AE, et al. (2014) A new tubulin-binding site and pharmacophore for microtubule-destabilizing anticancer drugs. Proc Natl Acad Sci U S A 111(38):13817–13821.

47. Takahashi M, et al. (1987) Rhizoxin binding to tubulin at the maytansine-binding site. Biochim Biophys Acta 926(3):215–223.

48. Partida-Martinez LP, et al. (2007) Burkholderia rhizoxinica sp. nov. and Burkholderia endofungorum sp. nov., bacterial endosymbionts of the plant-pathogenic fungus Rhizopus microsporus. Int J Syst Evol Microbiol 57(Pt 11):2583–2590.

49. Zhang L, Morrison M, & Rickard CM (2014) Draft Genome Sequence of Ralstonia pickettii AU12-08, Isolated from an Intravascular Catheter in Australia. Genome Announc 2(1).

50. Chen YY, et al. (2017) An Outbreak of Ralstonia pickettii Bloodstream Infection Associated with an Intrinsically Contaminated Normal Saline Solution. Infect Control Hosp Epidemiol 38(4):444–448.

51. Tejera D, Limongi G, Bertullo M, & Cancela M (2016) Ralstonia pickettii bacteremia in hemodialysis patients: a report of two cases. Rev Bras Ter Intensiva 28(2):195–198.

52. Ryan MP & Adley CC (2013) The antibiotic susceptibility of water-based bacteria Ralstonia pickettii and Ralstonia insidiosa. J Med Microbiol 62(Pt 7):1025–1031.

53. Casadevall A, Fu MS, Guimaraes AJ, & Albuquerque P (2019) The ‘Amoeboid Predator-Fungal Animal Virulence’ Hypothesis. J Fungi (Basel) 5(1).

54. Dunn JD, et al. (2017) Eat Prey, Live: Dictyostelium discoideum As a Model for Cell-Autonomous Defenses. Front Immunol 8:1906.

55. Cosson P & Soldati T (2008) Eat, kill or die: when amoeba meets bacteria. Current opinion in microbiology 11(3):271–276.

56. Hillmann F, et al. (2015) Virulence determinants of the human pathogenic fungus Aspergillus fumigatus protect against soil amoeba predation. Environ Microbiol 17(8):2858–2869.

57. Watkins RA, et al. (2018) Cryptococcus neoformans Escape From Dictyostelium Amoeba by Both WASH-Mediated Constitutive Exocytosis and Vomocytosis. Front Cell Infect Microbiol 8:108.

58. Koller B, et al. (2016) Dictyostelium discoideum as a Novel Host System to Study the Interaction between Phagocytes and Yeasts. Front Microbiol 7:1665.

59. King JS & Kay RR (2019) The origins and evolution of macropinocytosis. Philos Trans R Soc Lond B Biol Sci 374(1765):20180158.

60. Benghezal M, et al. (2006) Specific host genes required for the killing of Klebsiella bacteria by phagocytes. Cell Microbiol 8(1):139–148.

61. Partida-Martinez LP & Hertweck C (2007) A gene cluster encoding rhizoxin biosynthesis in “Burkholderia rhizoxina”, the bacterial endosymbiont of the fungus Rhizopus microsporus. Chembiochem: a European journal of chemical biology 8(1):41–45.

62. Mobius N & Hertweck C (2009) Fungal phytotoxins as mediators of virulence. Curr Opin Plant Biol 12(4):390–398.

63. Blin K, et al. (2017) antiSMASH 4.0-improvements in chemistry prediction and gene cluster boundary identification. Nucleic Acids Res 45(W1):W36–W41.

64. Scherlach K, Partida-Martinez LP, Dahse HM, & Hertweck C (2006) Antimitotic rhizoxin derivatives from a cultured bacterial endosymbiont of the rice pathogenic fungus Rhizopus microsporus. J Am Chem Soc 128(35):11529–11536.

65. Lackner G, Moebius N, & Hertweck C (2011) Endofungal bacterium controls its host by an hrp type III secretion system. ISME J 5(2):252–261.

66. Renshaw SA, et al. (2006) A transgenic zebrafish model of neutrophilic inflammation. Blood 108(13):3976–3978.

67. Ellett F, Pase L, Hayman JW, Andrianopoulos A, & Lieschke GJ (2011) mpeg1 promoter transgenes direct macrophage-lineage expression in zebrafish. Blood 117(4):e49–56.

68. Moebius N, Uzum Z, Dijksterhuis J, Lackner G, & Hertweck C (2014) Active invasion of bacteria into living fungal cells. eLife 3:e03007.

69. Shaffer JP, U’Ren JM, Gallery RE, Baltrus DA, & Arnold AE (2017) An Endohyphal Bacterium (Chitinophaga, Bacteroidetes) Alters Carbon Source Use by Fusarium keratoplasticum (F. solani Species Complex, Nectriaceae). Front Microbiol 8:350.

70. Hoffman MT, Gunatilaka MK, Wijeratne K, Gunatilaka L, & Arnold AE (2013) Endohyphal bacterium enhances production of indole-3-acetic acid by a foliar fungal endophyte. PLoS One 8(9):e73132.

71. Araldi-Brondolo SJ, et al. (2017) Bacterial Endosymbionts: Master Modulators of Fungal Phenotypes. Microbiol Spectr 5(5).

72. Uehling J, et al. (2017) Comparative genomics of Mortierella elongata and its bacterial endosymbiont Mycoavidus cysteinexigens. Environ Microbiol 19(8):2964–2983.

73. Takashima Y, et al. (2018) Prevalence and Intra-Family Phylogenetic Divergence of Burkholderiaceae-Related Endobacteria Associated with Species of Mortierella. Microbes Environ.

74. Gee JE, et al. (2011) Characterization of Burkholderia rhizoxinica and B. endofungorum isolated from clinical specimens. PLoS One 6(1):e15731.

75. Nazir R, Tazetdinova DI, & van Elsas JD (2014) Burkholderia terrae BS001 migrates proficiently with diverse fungal hosts through soil and provides protection from antifungal agents. Front Microbiol 5:598.

76. Frey-Klett P, et al. (2011) Bacterial-fungal interactions: hyphens between agricultural, clinical, environmental, and food microbiologists. Microbiology and molecular biology reviews: MMBR 75(4):583–609.

77. Benoit I, et al. (2015) Bacillus subtilis attachment to Aspergillus niger hyphae results in mutually altered metabolism. Environ Microbiol 17(6):2099–2113.

78. Zheng H, Dietrich C, & Brune A (2017) Genome Analysis of Endomicrobium proavitum Suggests Loss and Gain of Relevant Functions during the Evolution of Intracellular Symbionts. Appl Environ Microbiol 83(17).

79. Salvioli A, et al. (2016) Symbiosis with an endobacterium increases the fitness of a mycorrhizal fungus, raising its bioenergetic potential. Isme j 10(1):130–144.

80. Lackner G, Partida-Martinez LP, & Hertweck C (2009) Endofungal bacteria as producers of mycotoxins. Trends in microbiology 17(12):570–576.

81. Sharmin D, et al. (2018) Comparative Genomic Insights into Endofungal Lifestyles of Two Bacterial Endosymbionts, Mycoavidus cysteinexigens and Burkholderia rhizoxinica. Microbes Environ 33(1):66–76.

82. Sephton-Clark P, et al. (2019) Host-pathogen transcriptomics of macrophages, Mucorales and their endosymbionts: a polymicrobial pas de trois. BioRXiv.

83. Horn M & Wagner M (2004) Bacterial endosymbionts of free-living amoebae. J Eukaryot Microbiol 51(5):509–514.

84. Brock DA, Douglas TE, Queller DC, & Strassmann JE (2011) Primitive agriculture in a social amoeba. Nature 469(7330):393–396.

85. Brock DA, et al. (2020) Endosymbiotic adaptations in three new bacterial species associated with Dictyostelium discoideum: Paraburkholderia agricolaris sp. nov., Paraburkholderia hayleyella sp. nov., and Paraburkholderia bonniea sp. nov. PeerJ 8:e9151.

86. DiSalvo S, et al. (2015) Burkholderia bacteria infectiously induce the proto-farming symbiosis of Dictyostelium amoebae and food bacteria. Proc Natl Acad Sci U S A 112(36):E5029–5037.

87. Garcia JR, Larsen TJ, Queller DC, & Strassmann JE (2019) Fitness costs and benefits vary for two facultative Burkholderia symbionts of the social amoeba, Dictyostelium discoideum. Ecol Evol 9(17):9878–9890.

88. Stallforth P, et al. (2013) A bacterial symbiont is converted from an inedible producer of beneficial molecules into food by a single mutation in the gacA gene. Proc Natl Acad Sci U S A 110(36):14528–14533.

89. Boulais J, et al. (2010) Molecular characterization of the evolution of phagosomes. Mol Syst Biol 6:423.

90. Casadevall A (2012) Amoeba provide insight into the origin of virulence in pathogenic fungi. Adv Exp Med Biol 710:1–10.

91. Kontoyiannis DP, Wessel VC, Bodey GP, & Rolston KV (2000) Zygomycosis in the 1990s in a tertiary-care cancer center. Clinical infectious diseases: an official publication of the Infectious Diseases Society of America 30(6):851–856.

92. King JS, Veltman DM, Georgiou M, Baum B, & Insall RH (2010) SCAR/WAVE is activated at mitosis and drives myosin-independent cytokinesis. J Cell Sci 123(Pt 13):2246–2255.

93. Schindelin J, et al. (2012) Fiji: an open-source platform for biological-image analysis. Nat Methods 9(7):676–682.

94. Brothers KM, Newman ZR, & Wheeler RT (2011) Live imaging of disseminated candidiasis in zebrafish reveals role of phagocyte oxidase in limiting filamentous growth. Eukaryot Cell 10(7):932–944.

95. Kimmel CB, Ballard WW, Kimmel SR, Ullmann B, & Schilling TF (1995) Stages of embryonic development of the zebrafish. Developmental dynamics: an official publication of the American Association of Anatomists 203(3):253–310.

