## Supplemental Figure S1 for "Environmental interactions with amoebae as drivers of bacterial-fungal endosymbiosis and pathogenicity"

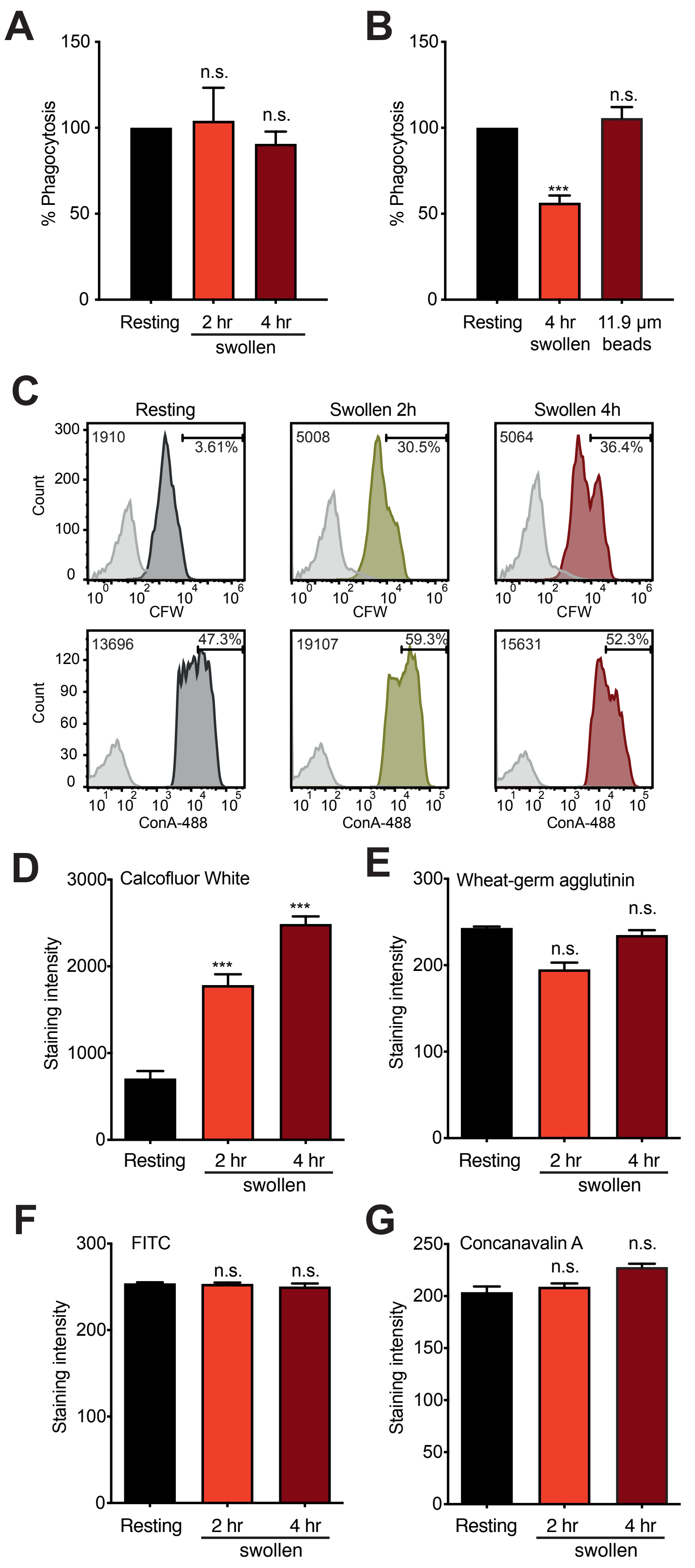


**Supplementary Figure 1**: Effect of spore viability and changes in cell wall composition upon swelling. (A) Phagocytosis of UV-inactivated resting and R. microsporus FP 469-12 spores by J774A.1 macrophages. (B) Phagocytosis of swollen (4h, 6.5  m) R. microsporus FP 469-12 spores and latex beads (11.9  m) by J774A.1 macrophages. For both experiments the number of phagocytes containing at least one spore or bead were scored as positive. (C) Total Chitin (CFW) and Mannan (ConA) exposure in resting and swollen spores analysed by flow cytometry (>10,000 cells), including Median Fluorescence Intensity (left) and Percent High (right) for each population. Data representative of three independent repeats is shown. (D-E) Shows quantification of changes in cell wall staining upon swelling of parental R. microsporus FP 469-12 spores, visualized by fluorescence microscopy. (D) Shows total chitin (Calcofluor White staining), (E) exposed chitin (Wheat-Germ Agglutinin), (F) total protein (FITC) and (G) mannan (Concanavalin A). All graphs show mean ± SEM of 3 independent repeats. ns = p>0.05, ***=p<0.0001, One-Way ANOVA with Tukey's correction for multiple comparisons.
