## Supplemental figure S2 for "Environmental interactions with amoebae as drivers of bacterial-fungal endosymbiosis and pathogenicity"

**A**

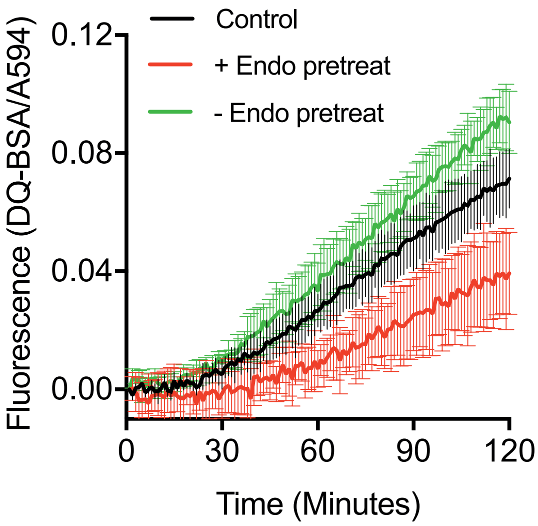


**Supplementary figure 2:** Pretreatment of D. discoideum with conditioned medium has no additional effect. (A) D. discoideum cells were pretreated with medium conditioned by R. microsporus FP 469-12 spores with and without endosymbionts for 30 minutes prior to addition of DQ-BSA proteolysis reporter beads.
