## Supplemental Figure S3 for "Environmental interactions with amoebae as drivers of bacterial-fungal endosymbiosis and pathogenicity"

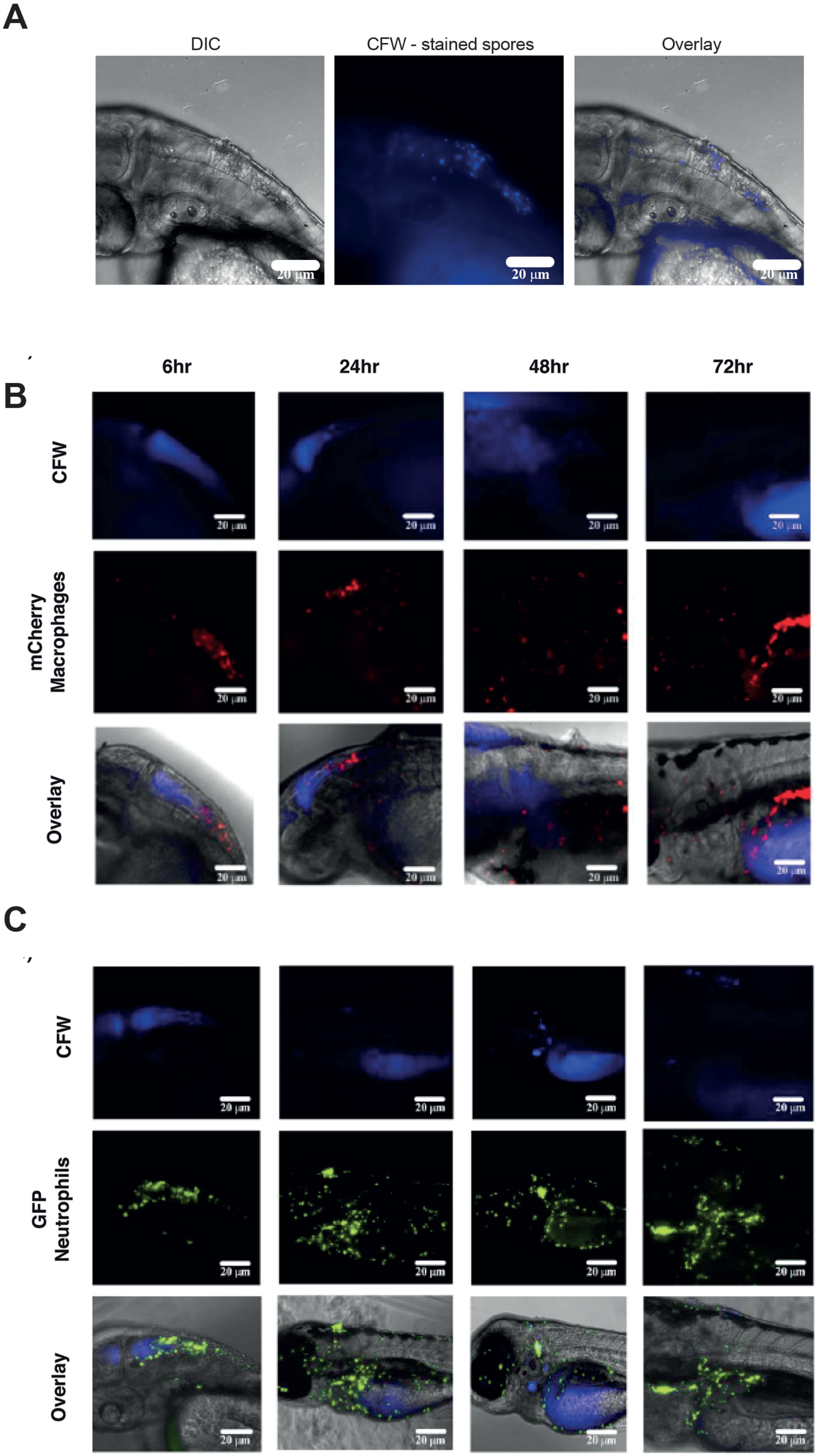


**Supplementary Figure 3**: Zebrafish larval infection model. (A) Representative micrographs of zebrafish larvae infected with CFW-stained swollen parent spores as analysed in Figure 7. (B) Representative images of macrophage-reporter zebrafish (mpeg1:G/U:NfsB-mCherry) showing recruitment to sites of infection with CFW-stained fungal spores. (C) Equivalent images of infected GFP-neutrophil (mpx:GFP)i114)-reporter fish.
