## Supplemental Figure S4 for "Environmental interactions with amoebae as drivers of bacterial-fungal endosymbiosis and pathogenicity"

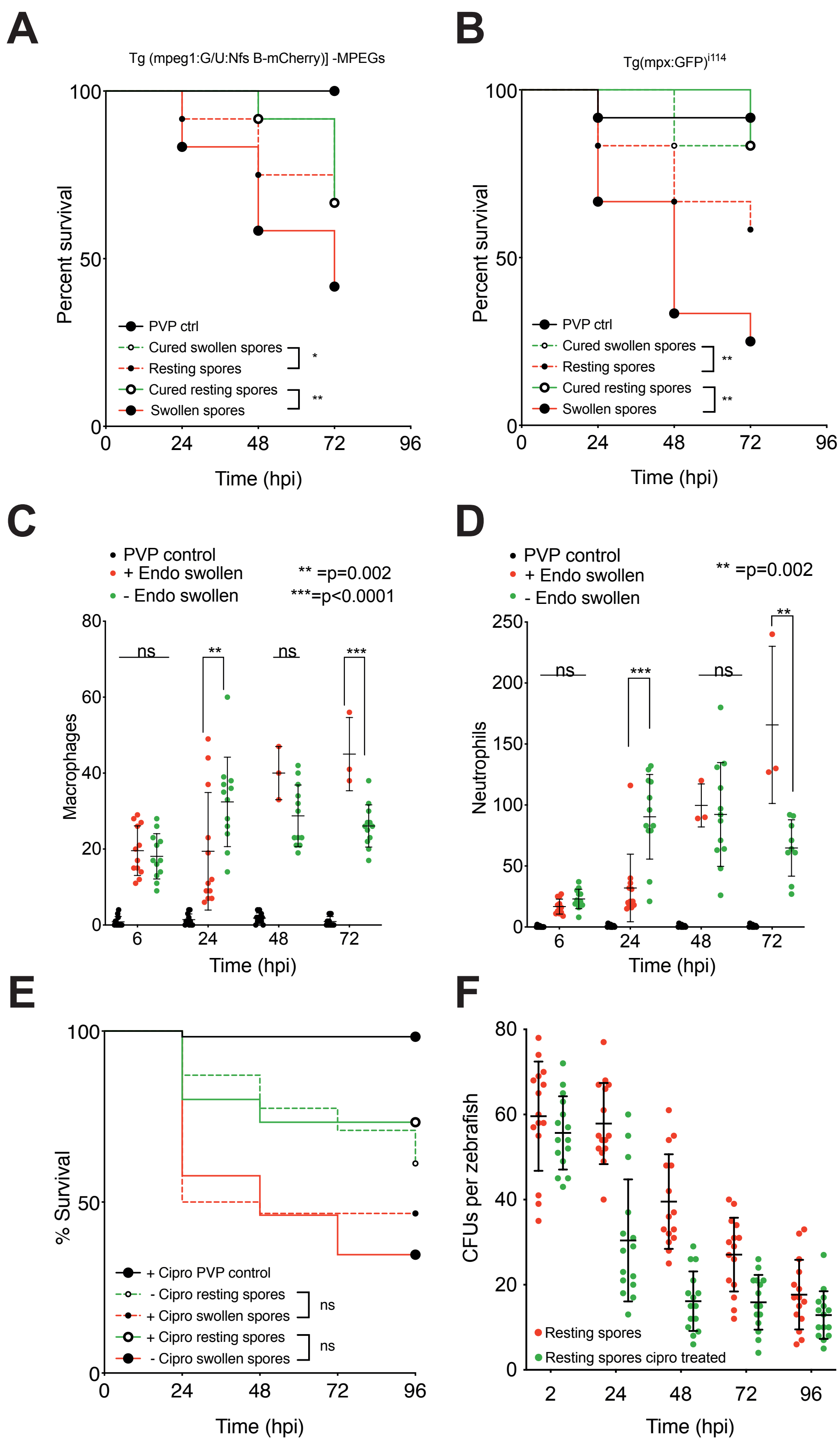


**Supplementary figure 4:** Effect of endosymbiont status on infection of zebrafish with swollen spores. (A) Kaplan-Meijer survival curves of transgenic macrophage-reporter (Tg(mpeg1:G/U:NfsB-mCherry)) zebrafish following hindbrain injection with R. microsporus FP 469-12 spores with and without endosymbionts. (B) Equivalent survival data of neutrophil-reporter (Tg(mpx-:GFP)i114) zebrafish. n=12 fish per condition, statistical differences were determined using Mantel-Cox with Bonferroni's correction for multiple comparisons (5% family-wise significance threshold = 0.025). (C) and (D) Effect of infection with pre-swollen spores (4hr) on macrophage and neutrophil recruitment respectively. Three biological replicates of 4 fish per condition were examined (n=12). Statistical significance was assessed by Two-way ANOVA with Tukey's correction for multiple comparisons or pairwise t-tests where sample number was unequal due to fish death. (E) Survival curves for impact of 60 g/ml ciprofloxacin on survival of wildtype AB fish upon hindbrain infection with resting or swollen parental R. microsporus FP 469-12 spores (n=30). Injected fish were then cultivated in E3 medium with or without 60 g/ml Ciprofloxacin and monitored for mortality (n=30). Statistical differences were determined using Mantel-Cox with Bonferroni's correction for multiple comparisons (5% family-wise significance threshold = 0.025). (F) Shows CFU's recovered from fish treated as in (E). No statistical significance was observed at any time point. In all panels *=p<0.05; **=p<0.001; ***=p<0.0001 unless otherwise indicated.
